# Microenvironmental Ammonia Enhances T cell Exhaustion in Colorectal Cancer

**DOI:** 10.1101/2022.05.25.493422

**Authors:** Hannah N. Bell, Amanda K. Huber, Rashi Singhal, Ryan J. Rebernick, Roshan Kumar, Nupur K. Das, Samuel A. Kerk, Peter Sajjakulnukit, Sumeet Solanki, Jadyn G. James, Donghwan Kim, Li Zhang, Marwa O. El-derany, Timothy L. Frankel, Balázs Győrffy, Eric R. Fearon, Marina Pasca di Magliano, Frank J. Gonzalez, Ruma Banerjee, Costas A. Lyssiotis, Michael Green, Yatrik M. Shah

## Abstract

Effective therapies are lacking for patients with advanced colorectal cancer (CRC). The CRC tumor microenvironment has elevated metabolic waste products due to altered metabolism and proximity to the microbiota. The role of metabolite waste in tumor development, progression, and treatment resistance is unclear. We generated an autochthonous metastatic mouse model of CRC and unbiased multi-omic analyses in this model reveals a robust accumulation of tumoral ammonia. The high ammonia levels induce T cell metabolic reprogramming, increase exhaustion and decrease proliferation. CRC patients have increased serum ammonia, and our ammonia-related gene signature correlates with altered T cell response, adverse patient outcomes, and lack of response to immune checkpoint blockade. We demonstrate that enhancing ammonia clearance reactivates T cells, decreases tumor growth, and extends survival. Moreover, decreasing tumor-associated ammonia enhances anti-PD-L1 efficacy. Our findings indicate that ammonia detoxification can reactivate T cells, highlighting a new approach to enhance the efficacy of immunotherapies.

**Statement of Significance:** We demonstrate that ammonia accumulates in the microenvironment of colorectal cancer. Ammonia alters T-cells redox singling leading to a decrease in T cell proliferation and an increase in T cell exhaustion. Enhancing ammonia clearance reduces tumor size, increases survival, and increases the efficacy to immunotherapies.

## INTRODUCTION

Colorectal cancer (CRC) is the second leading cause of cancer mortality worldwide(1). Improvements in CRC patient survival stems in part from enhanced early detection due to increased rates of colonoscopy (2). However, 20% of patients with CRCs have metastatic disease at the time of diagnosis, and the 5-year overall survival rate in this patient subset is 10.5% (3,4). Moreover, immune checkpoint blockade (ICB) is not effective in the vast majority of patients with CRC (5). Recent work has revealed that the tumor microenvironment (TME) in many CRCs is often immunosuppressive. The TME is a hypoxic, nutrient-sparse environment with high concentrations of metabolic waste. Heterocellular metabolite exchange in the TME can lead to dysregulated immune cell effector function (Beckermann et al., 2017; Boedtkjer & Pedersen, 2019; Hanahan, 2022). Metabolic dysregulation in primary CRCs is also affected by intestinal microbes, as the gut microbiota generates unique metabolites and waste products.

Ammonia is a waste product that is generated at high levels from host and microbial cellular metabolism. Host-derived ammonia arises via glutamine and asparagine catabolism and cysteine and nucleotide biosynthesis(6,7). Microbiota also produce large quantities of ammonia through breakdown of host proteins by microbial ureases (8). Due to its toxic effects on many cells, ammonia must be exported and processed by the liver through the urea cycle (9). Of note, some cancer cells can assimilate ammonia as a biosynthetic metabolite to drive amino acid metabolism (10). However, the role of extracellular ammonia in the TME is largely unstudied.

We employed an autochthonous metastatic model of CRC arising from concurrent mutations in tumor suppressor and oncogenes known to contribute to human CRC. Using multiomic profiling to investigate the altered immune landscape and mechanisms of immune therapy resistance, we identified an important transcriptional network regulated by hepatic nuclear factor-4 (HNF4)α. HNF4α is a master regulator of genes encoding urea cycle factors, and disruption of HNF4α leads to increased ammonia accumulation (11). We identify a robust increase of ammonia in a metastatic colon cancer model and show increased TME ammonia levels reduce T cell activation and proliferation and lead to T cell exhaustion. Promoting enhanced TME ammonia detoxification inhibits tumor growth and activates the response to ICB in our mouse CRC model. Evidence implicates elevated ammonia levels in the TME of human CRC patients that do not respond to ICB and/or who have poorer outcomes. Collectively, our findings identify ammonia as a key immune-inhibitory waste product in CRC and highlight the potential of enhancing ammonia clearance as an approach to increase the effectiveness of ICB in CRC.

## RESULTS

### Mouse CRC model lacks response to multiple immune therapy approaches

ICB therapy has recently led to clinical breakthroughs for some solid tumors, such as clear cell renal cell carcinoma, melanoma, non-small cell lung cancer, and mismatch repair defective (MMRD) CRCs and other MMRD cancers(12,13). However, in the vast majority of CRCs, ICB is not effective (Diao et al., 2020). Unlike the situation in human CRC, some widely used syngeneic mouse CRC xenograft models as well as sporadic mouse CRC models are highly responsive to immunotherapies (Efremova et al., 2018). To understand how genetically engineered mouse models with variable defects in the murine homologues of tumor suppressor and oncogenes recurrently mutated in human CRC would respond to immune-based therapies, we pursued work with mice where tamoxifen treatment can induce target genes of interest in mouse colon epithelium to induce benign or malignant tumors (Bell et al., 2021; Feng et al., 2011). Mice with a biallelic disruption of *Apc* (*SingleMut*), *Apc* and *Tp53* (*DoubleMut*), or *Apc, Tp53* and a *Kras* G12D knock-in allele (*TripleMut*) were assessed for overall survival following one, two, three, or four daily doses of 100 mg/kg tamoxifen (**Figure 1A and B, S1A**). As expected, increasing tamoxifen doses led to progressively shorter survival times. Of note, for equivalent tamoxifen dosages, the average survival of the *TripleMut* is approximately 50% shorter than that of the *DoubleMut*. Moreover, *TripleMut* had increased numbers of tumors, and histologically more advanced tumors compared to *SingleMut* or *DoubleMut* (**Figure 1C, 1D, 1E**). The *TripleMut*, but not the *SingleMut* and *DoubleMut* mice developed lung and liver metastases (**Figure 1F)**. We assessed the response of each mouse model to anti-Programmed-Dealth-Ligand-1 (PD-L1). *SingleMut and DoubleMut* demonstrated increased survival following anti-PD-L1 treatment (**Figure 1G**). However, *TripleMut* did not show improved survival following anti-PD-L1 treatment as compared to isotype control (**Figure 1G**). We next used an alternative form of immunotherapy, IL-17 bone marrow knockout, in the *TripleMut* mouse model. IL-17 promotes colorectal cancer by driving pro-tumor inflammation, generating an immunosuppressive microenvironment, and suppressing cytotoxic immune function (W.-J. Chae et al., 2010; Grivennikov et al., 2012; Wu et al., 2013). Using bone marrow chimeras from IL-17 knockout mice, we found that the *SingleMut* mice had increased survival following disruption of IL-17 signaling as previously reported, but the *TripleMut* mouse model showed no response to transplantation with IL-17-disrupted immune cells (**Figure S1B**) (W.-J. Chae et al., 2010).) Together, these data reveal that the *TripleMut* model is resistant to immune-based therapeutic approaches.

**Figure 1:**
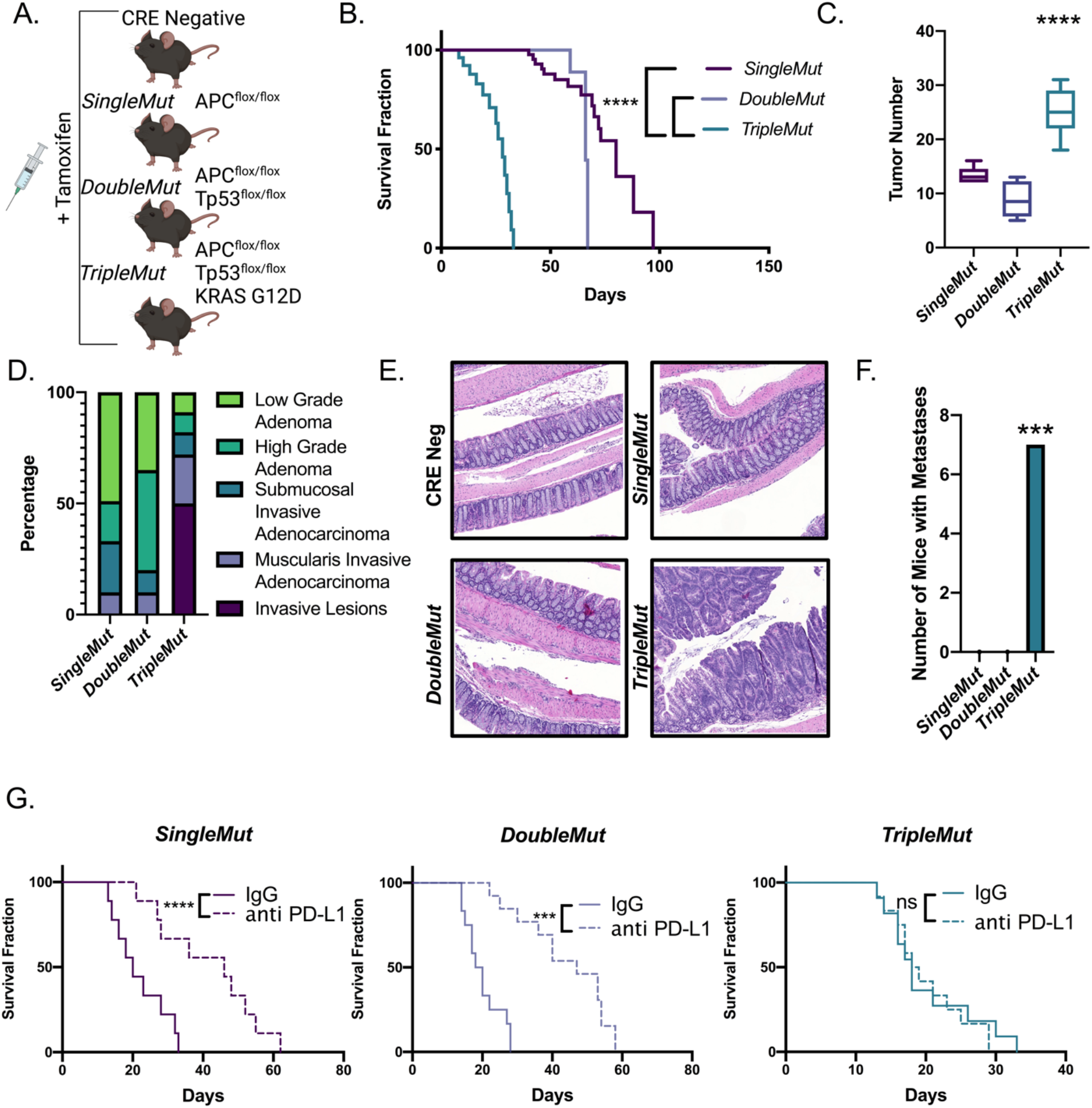
Autochthonous metastatic mouse model of colon cancer is resistant to immunotherapy. A) Schematic of *SingleMut, DoubleMut*, and *TripleMut*. B) Mice were induced with one injection of 50 mg/kg tamoxifen and then assessed for survival, C) tumor number (n=8-10), D) tumor progression, and E) histology (n=8-10). F) Mice were treated with low dose tamoxifen and lungs and livers were macroscopically assessed for metastasis. G) Mice were induced with tamoxifen, treated daily with intraperitoneal anti-PD-L1 (1 μg per mouse) and monitored for survival. (n=8-10). Statistics were calculated with one-way ANOVA or t-test. Survival significance is calculated using log-rank test. *p < 0.05, ** p < 0.01, *** p < 0.001,**** p < 0.0001 unless otherwise indicated. Data is presented as mean +/- the standard error of the mean.

### Mice with advanced lesions develop an immunosuppressive microenvironment

We sought to understand the differential responsiveness of the CRC models to immune-based therapies. Acquired and adaptive resistance to immunotherapy can result from dysfunction or deletion of CD8+ T cells. RNA sequencing on colon epithelium demonstrated differential global gene expression for each model **(Table S1)**. Principal components analyses demonstrated that the *TripleMut* mice separated from the *SingleMut* and *DoubleMut* mice along PC1 (**Figure 2A and B**). Gene set enrichment analysis revealed a robust depletion in pathways essential for adaptive immune response or T cell function in *TripleMut* mice compared to the *DoubleMut* (**Figure 2C**). We performed multiplex immunohistochemical staining (OPAL) for CD3, Arginase, CD8 and CK19 on paraffin sections from a different mouse cohort (**Figure S2A**). The *TripleMut* mouse model showed an increase in arginase positive cells (**Figure S2B**). We saw a decrease in the distance between CD3 positive T cells and arginase positive myeloid cells, suggesting possible contacts between immunosuppressive macrophages and tumorinfiltrating T cells (**Figure S2C**). We next performed CyToF on a separate cohort of mice (**Figure 2D**). *TripleMut* mice had increased CD44+/CD45-cells, indicating an increased proportion of cancer stem cells in this mouse model (**Figure 2E**). The *TripleMut* mouse model also displayed a significant decrease in CD45+ cells and CD3+ cells (**Figure 2E**). The number of CD4+ cells and CD8+ T cells in the *TripleMut* model was decreased, but not significantly (**Figure S2D**). We observed no significant changes in the CD11b+ Ly6G+ or CD11b+ F4/80+ populations between the *DoubleMut* and *TripleMut* mice (**Figure S2E**). The *TripleMut* mouse model had a significant decrease in activated CD4+ T cells and a significant increase in CD4+ and CD8+ positive T cells expressing the inhibitory receptors CTLA-4 and PD-1 (**Figure 2F and S2F**). Overall, the findings suggest the *TripleMut* mice have fewer overall T cells and increased T cell dysfunction.

**Figure 2:**
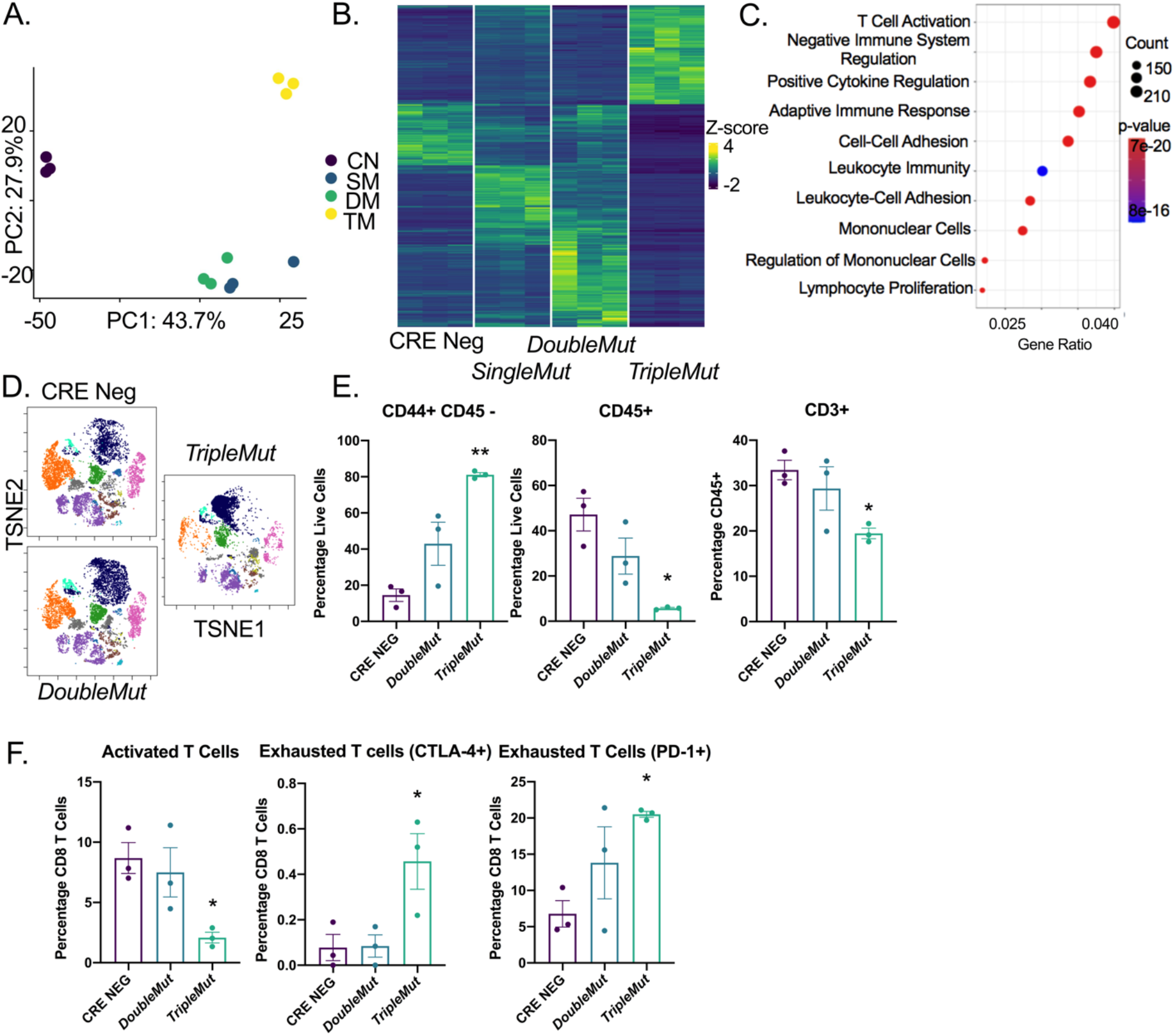
T cells are reduced in the *TripleMut* mice. A) PCA analysis of RNA-sequencing of colon cancer mouse models induced with one dose of 100 mg/kg tamoxifen (n=3). B) Heat map and C) gene set enrichment of differentially expressed genes between mouse models (n=3). D) Representative TSNE plot of CyToF data of different mouse models induced with one dose of 100 mg/kg tamoxifen (n=3). E and F) Selected cell populations from CyToF data (n=3). *p < 0.05, ** p < 0.01, *** p < 0.001, **** p < 0.0001 unless otherwise indicated. Data is presented as mean +/- the standard error of the mean.

### Ammonia accumulates in CRC due to dysregulation of the urea cycle

To analyze which transcription factors were responsible for the differential expression profile between the *DoubleMut* and *TripleMut* mice, a transcription factor enrichment analysis was performed (Keenan et al., 2019). We screened the enriched transcription factors for expression levels that correlated with survival, differential expression in CRC compared to normal colon, and an association with KRAS using STRING was assessed (Szklarczyk et al., 2018). This analysis identified four potential candidates: GFI, HNF4α, RUNX1, and the IRF protein family (**Figure 3A**). HNF4α is a master regulator of the urea cycle and is necessary to clear cellular ammonia, and *TripleMut* mice exhibited altered expression of genes critical for the generation or clearance of ammonia (*Gls1, Glud, Otc, Cps1*, and *Slc4a11*) (Inoue et al., 2002) (**Figure 3B and 3C**). Metabolomics analysis, using PCA analysis on *SingleMut*, *DoubleMut*, and *TripleMut* mice, found the metabolic profiles clustered separately (**Table S2, Figure S3A**). Urea cycle intermediates were not targeted with this particular metabolomics analysis, but differentially abundant metabolites in the *TripleMut* identified a robust decrease in pyrimidine metabolism (**Figure S3B and S3C**). Dysregulated pyrimidine metabolism is a central feature in urea cycle dysfunction, as nitrogen is diverted towards carbamoyl-phosphate synthetase 2, aspartate transcarbamylase, and dihydroorotase activation (Lee et al., 2018). Hepatic nuclear factor 4 (HNF4)α and its direct target the Ornithine Transcarbamylase (OTC), were significantly downregulated in CRC compared to normal colon tissues (**Figure 3D**). These findings were confirmed in an institutional cohort of CRC patients (**Figure 3E**). Further, analysis of epithelial and stromal compartments from the *SingleMut, DoubleMut*, and *TripleMut* mouse models demonstrated that *Otc* and *Hnf4a* levels were reduced in the tumor epithelium (**Figure 3F**). Enteroids generated from the *TripleMut* mice had the most significantly reduced *Hnf4a* and *Otc* expression compared to wildtype, *SingleMut*, and *DoubleMut* enteroids **(Figure S3D)**. Similarly, the normal human colon cell line NCM460 highly expressed *HNF4a* and *OTC* compared to CRC-derived cell lines HCT116 and the SW480 (**Figure S3E**). Moreover, analysis of data from the Cancer Cell Line Encyclopedia and TCGA found that cell lines or patients with a KRAS mutation had lower levels of *HNF4a* (**Figure S3F and S3G**). To determine the role of mutant KRAS in the dysregulation of *Hnf4a* and *Otc*, a colon-epithelial doxycycline inducible *Kras* mutant mouse model was generated (**Figure S3H**). Mice expressing the mutant *Kras* allele had an increased prevalence of dysplasia, tumor formation, and increased GFP *Kras* reporter expression in their colons (**Figure S3I**). In the mutant *Kras* mice, *Hnf4a* and *Otc* expression were decreased, although this decrease was not significant (**Figure S3J**). This suggests that the downregulation of *Hnf4a* and *Otc* is a combinatorial effect of *Apc* and *Tp53* loss as well as mutant *Kras* expression.

**Figure 3:**
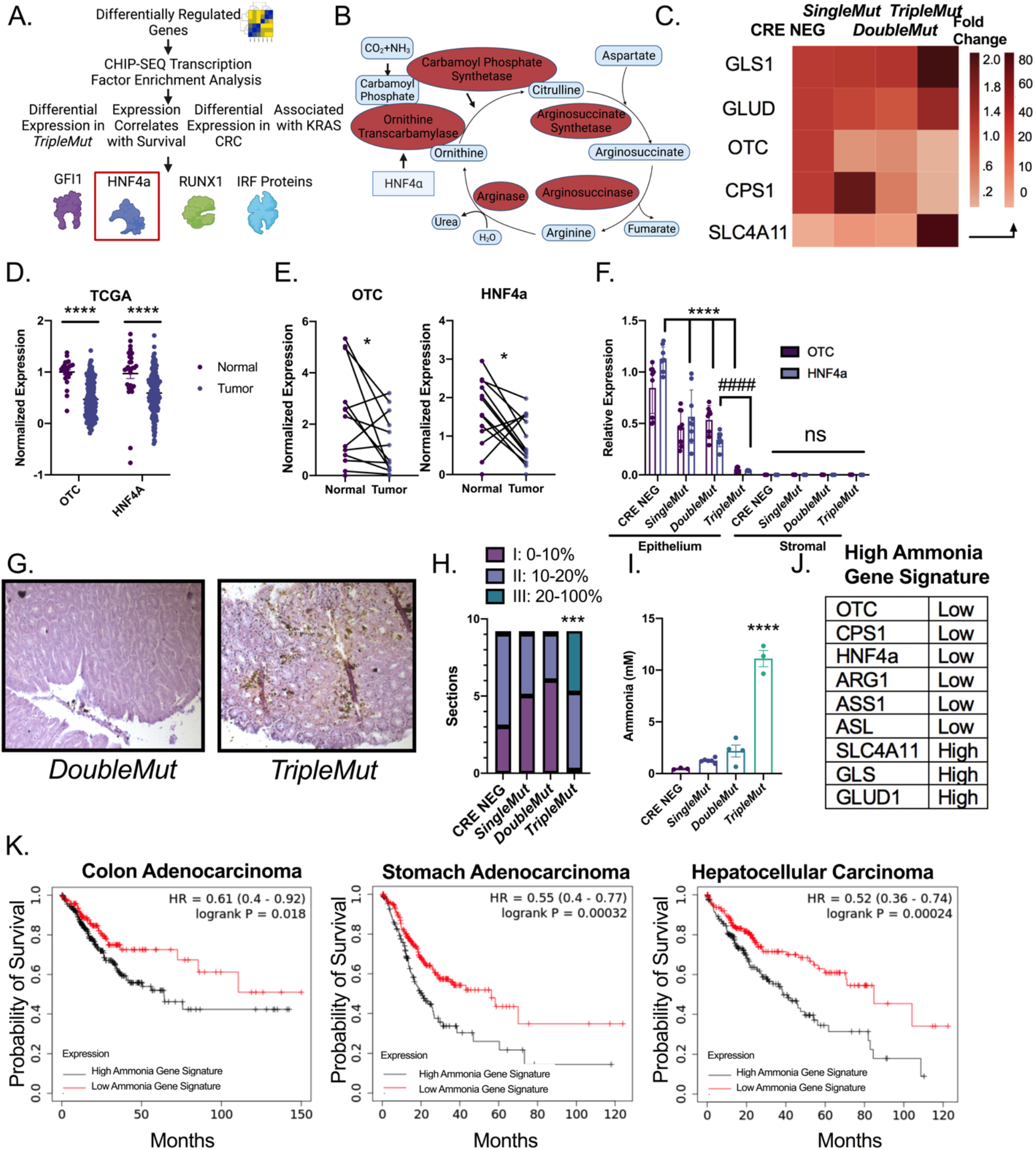
Ammonia metabolism is dysregulated in advanced colorectal cancer. A) Schematic of transcription factor analysis. B) Schematic of urea cycle intermediates and enzymes. C) Heat map of expression of genes (GLS1 (Glutaminase-1), GLUD (Glutamate Dehydrogenase-1), OTC (Ornithine Transcarbamylase), CPS1 (Carbamoyl Phosphate Synthase-1), and SLC4A11 (Solute Carrier Family 4 Member 11)) in the ammonia signature from RNA-Sequencing experiment. D) OTC and HNF4α expression in cancer and normal tissue from the TCGA dataset. E) Gene expression of HNF4α and OTC in normal and cancer patients. F) Gene expression of flow-sorted epithelial and stromal compartments. (n=6-8). G) Representative Nessler’s staining and H) quantification of mice induced with 100 mg/kg tamoxifen and sacrificed at endpoint (n=9). I) Mass spectrometry ammonia quantification of tumors from mice induced with 100 mg/kg tamoxifen. (n=3). J) Genes and expression comprising the high ammonia gene signature. K) Kaplan-Meier survival curves of cancer patients stratified by the ammonia gene signature. Data from TCGA. *p < 0.05, ** p < 0.01, *** p < 0.001, **** p < 0.0001 unless otherwise indicated. Data is presented as mean +/- the standard error of the mean.

The downregulation of *Hnf4a, Otc* and other ammonia related genes in our cancer models, and the metabolic shift in these metabolites prompted us to assess ammonia accumulation in CRC. Nessler’s reagent is a stain that can detect free ammonia in tissues (Zhao et al., 2019). Staining *DoubleMut* and *TripleMut* colons demonstrated that *TripleMut* mice had significantly increased epithelial ammonia staining compared to the *DoubleMut* (**Figure 3G and H**). To confirm these findings, we used mass spectrometry in an independent cohort of mice, and found a robust increase in ammonia in *TripleMut* mice (**Figure 3I**). Our ammonia gene signature consists of low expression of *OTC, CPS1, HNF4a, ARG1, ASS1*, and *ASL* and high expression of *SLC4A11, GLS*, and *GLUD1* (**Figure 3J**). In three gastrointestinal carcinomas (colon, gastric, and hepatocellular) the enrichment genes correlative to ammonia accumulation were associated with decreased survival (**Figure 3K**) (Lánczky & Győrffy, 2021). However, the signature did not predict survival in most other cancer types, potentially suggesting a specificity for the gastrointestinal tract. Together, these data reveal: that 1) dysregulation of ammonia metabolism in murine and human CRC leads to intratumoral ammonia accumulation, and 2) that ammonia-related gene expression prognostically stratifies CRC patients.

### Ammonia exposure increases exhausted T cells in vitro and in vivo

We assessed how increased microenvironmental ammonia altered the growth of a panel of CRC cell lines and primary immune cells. Ammonia increased or did not change cell growth in several CRC cell lines, consistent with what was previously reported in breast cancer-derived cell lines (Spinelli, Yoon, et al., 2017) (**Figure 4A**). While bone marrow derived neutrophils and macrophages were unaffected by ammonia levels, *in vitro* activated splenic CD3+ T cells showed a strong reduction in proliferation at doses within the range of ammonia observed in colon tumors. We confirmed that MC38 CRC cancer cell line and primary hepatocytes could clear ammonia following treatment, ammonia accumulates within T cells (**Figure S4A**). Dose response studies demonstrated significantly diminished proliferation of CD4+ and CD8+ T cells (**Figure 4B and S4B-C**), reduced expression of the T cell activation marker CD25 (**Figure 4C**), and a significant increase in cell death after ammonia treatment (**Figure 4D**). Further, human T cells treated *in vitro* with ammonia had significantly reduced proliferation (**Figure 4E**). Similar to the *TripleMut* mice, ammonia treatment increased the exhausted T cell markers (PD1+TIM3+), and decreased production of interferon gamma (IFNγ) in T cells as assessed by flow cytometry (**Figure 4F and S4D**). To confirm that ammonia drives an exhausted tumor microenvironment, we analyzed gene expression of PD-1 and CTLA-4 after ammonia treatment (**Figure 4G**). Consistent with the mouse data, ammonia significantly induced PD-1 expression, but not CTLA-4 in human T cells (**Figure 4H and S4E**). To determine whether an increase in systemic ammonia levels in mice alters T cell function, mice were placed on an ammonium acetate diet. After seven days, serum ammonia levels were increased (**Figure 4I**). T cells isolated from mice on ammonium acetate diet demonstrated decreased proliferation, as well as increased PD-1 and CTLA-4 expression, supporting the model that high levels of ammonia drive T cell exhaustion (**Figure 4J and 4K**). Interestingly, T cells treated with ammonia for 12 h, then replated with fresh media for 72 h, had a sustained decrease in T cell growth (**Figure S4F**). To confirm the long-term effects of ammonia on T cells, we treated mice with ammonia acetate or normal chow, isolated CD8+ T cells and adoptively transferred the cells into MC38 tumor bearing mice following *ex vivo* activation (**Figure 4L**). As expected, activated CD8+ T cells from control mice were able to reduce tumor growth, but T cells isolated from mice on the ammonium acetate chow did not suppress tumor growth (**Figure 4M**). Collectively, this data suggests that ammonia induces T cell exhaustion.

**Figure 4:**
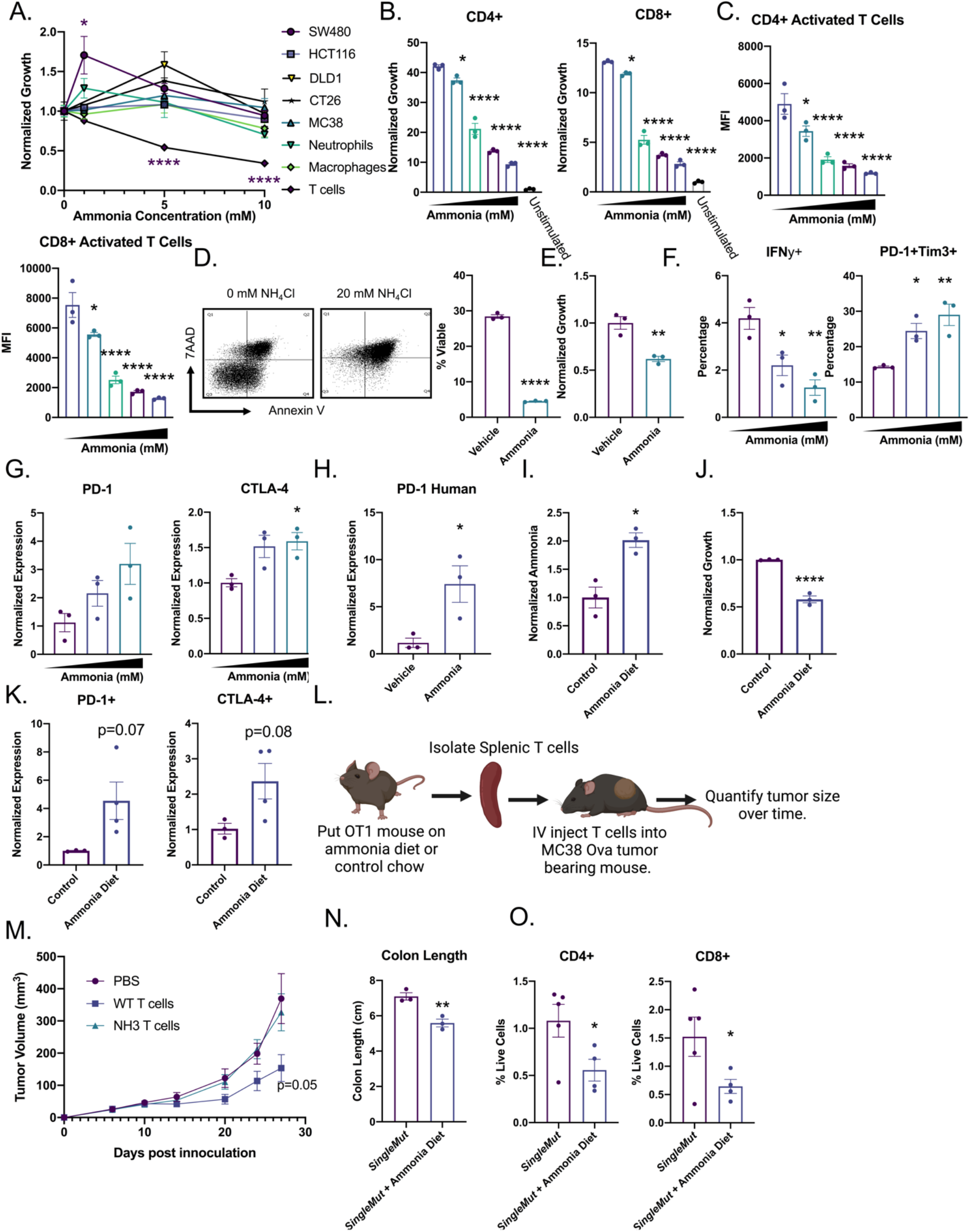
Ammonia increases exhausted T cells and reduces T cell viability. A) Panel of cell types treated with indicated ammonia concentration for 72 h. T cells were stimulated with CD3 and CD28. Quantification of T cell B) proliferation (CFSE), C) activation (CD25), and D) cell death (7AAD and Annexin V). Ammonia levels range from 1-10 mM. E) Proliferation of human T cells treated with ammonia for 72 h. F and G) IFNγ and exhaustion markers in mouse T cells by flow and RT-PCR after the indicated ammonia concentration for 72 h. H) Gene expression of human T cells treated with ammonia for 72 h. I) Serum ammonia quantification in mice treated for 7 days with ammonium acetate diet (n=3). J) Proliferation and K) gene expression of CD3 positive T cells isolated from the spleens of mice treated with ammonium acetate diet or control diet (n=3). L) Schematic illustrating the T cell transfer experiment in which mice were treated with control chow or ammonium acetate chow for 7 days. T cells were isolated, counted, and intravenously injected into MC38 tumor bearing mice. M) Tumors were monitored every 3 days (n=5-7). *SingleMut* mice were induced with 3 injections of 100 mg/kg tamoxifen and placed on ammonium acetate diet. At end point, N) mouse colon length O) intratumoral T cells were assessed (n=4). *p < 0.05, ** p < 0.01, *** p < 0.001, **** p < 0.0001 unless otherwise indicated. Data is presented as mean +/- the standard error of the mean. All experiments were performed in triplicates at least three times.

To understand if ammonia can drive tumorigenesis and alter tumor T cell response, an ammonium acetate diet was fed to the *SingleMut* mouse model. *SingleMut* mice on ammonium acetate compared to untreated *SingleMut* mice had decreased body weight, decreased survival, shorter and more dysplastic colons, and decreased proportion of CD4+ and CD8+ T cells (**Figure 4N and 4O and S4G**). This data indicates that ammonia treatment durably reduces the proliferation and activation of T cells and contributes to tumor progression.

### Ammonia inhibits the transsulfuration pathway to induce oxidative stress in T cells

To understand the mechanisms by which ammonia alters T cell function, targeted metabolomics was performed. T cells isolated from wildtype mice and treated *in vitro* with ammonia, and T cells isolated from mice on an ammonium acetate diet clustered on PC1, indicating a similar metabolic response (**Figure 5A, Table S3**). Amino acids, urea cycle and nicotinamide metabolites were the most enriched and significantly altered metabolites that were common between both treatment paradigms (**Figure 5B-C**)

**Figure 5:**
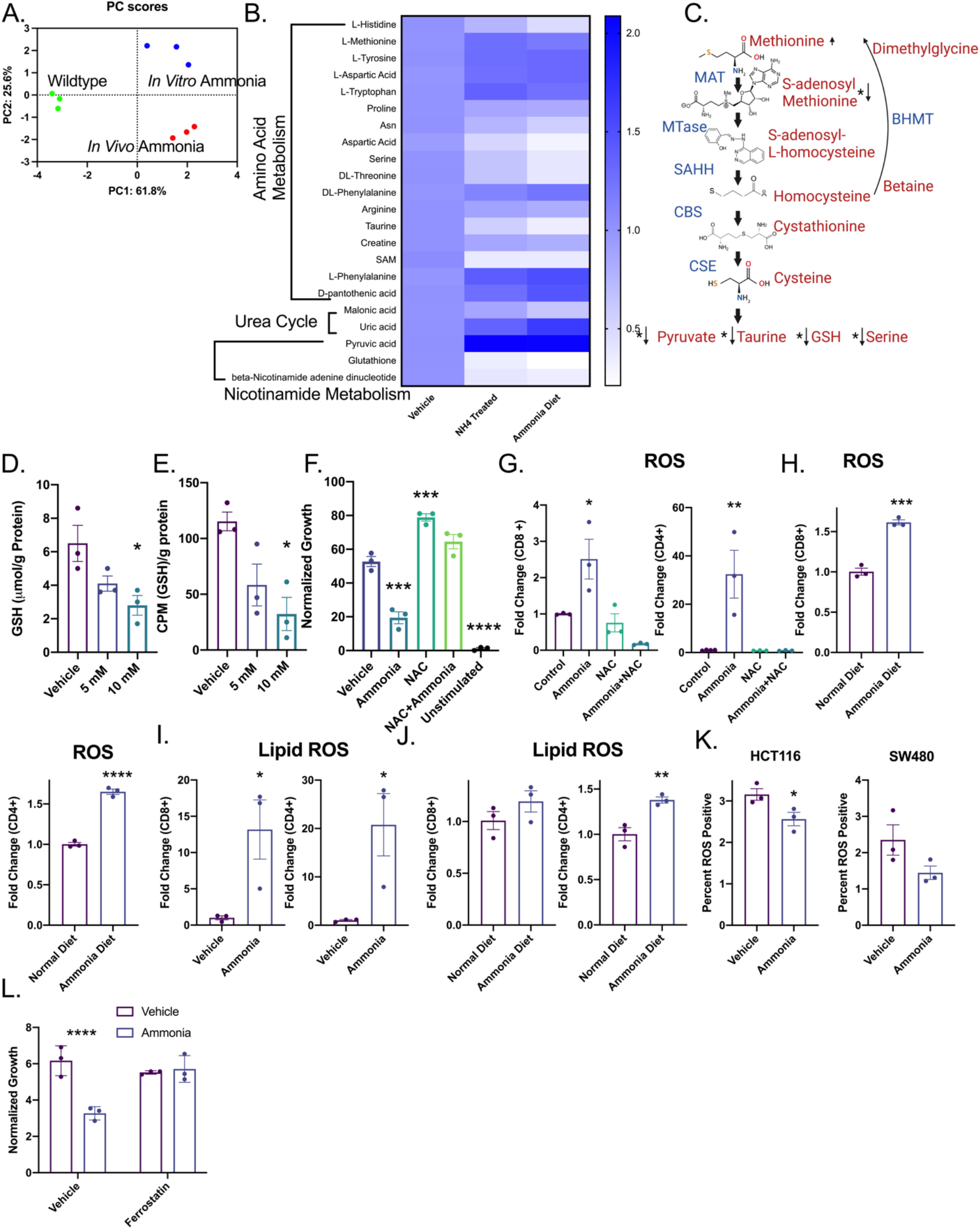
Ammonia leads to oxidative stress and dysregulated methionine metabolism in T cells. A) PCA analysis and B) heat maps annotated with significant pathways from pathway analysis of metabolomics in T cells isolated from wildtype mice, mice treated with ammonium acetate diet or wildtype T cells treated with 10 mM ammonia *in vitro* for 24 h (n=3). C) Schematic of the transsulfuration pathway with selected metabolite relative levels indicated. D) ^35^S tracing of methionine flux into GSH in *in vitro* treated mouse T cells. E) Absolute quantification of GSH in *in vitro* treated mouse T cells. F) T cells were treated for 24 h with 10 mM NAC and then treated with 10 mM ammonia for 72 h before CFSE was acquired using flow cytometry. G) T cells were treated with ammonia or cotreated with ammonia and NAC, and ROS was assessed by flow cytometry. H) Mice were treated with ammonium acetate diet for 7 days, then splenic T cells were assessed for ROS by flow cytometry (n=3). I) T cells were treated with ammonia and lipid ROS was assessed by flow cytometry. J) Mice were treated with ammonium acetate diet for 7 days, and isolated T cells were assessed for lipid ROS by flow cytometry (n=3). K) CRC-derived cell lines were treated with 10 mM ammonia for 72 h, assessed for lipid ROS by flow cytometry. L) T cells were pretreated for 24 h with ferrostatin (2 μm) followed by cotreatment with 10 mM ammonia for 24 h. *p < 0.05, ** p < 0.01, *** p < 0.001, **** p < 0.0001 unless otherwise indicated. Data is presented as mean +/- the standard error of the mean. All cell experiments were performed in triplicates at least three times.

Notably, there were robust changes in the transsulfuration pathway (**Figure 5C**). To evaluate the dysregulation of the transsulfuration pathway, total protein methylation was assessed. Histone methylation was not affected in T cells treated with ammonia *in vitro* or *in vivo* (**Figure S5A and S5B**). CBS and MAT2A expression were also unchanged (**Figure S5A**). However, gene expression of enzymes in the transsulfuration pathway identified betaine-homocysteine S-methyltransferase (BHMT) as an ammonia regulated gene in T cells **(Figure S5C and S5D)**. *Bhmt* was unchanged in ammonia treated MC38 colorectal cancer cells (**Figure S5E**). Since glutathione was significantly reduced following ammonia treatment **(Figure 5D)**, a radiolabel flux analysis was performed using [35S]-methionine to directly assess if defects in the transsulfuration pathway led to a decrease in glutathione. Radiolabel incorporation into glutathione was significantly decreased following ammonia treatment **(Figure 5E)**. While proliferation was not rescued by supplementing with additional amino acids or S-adenosylmethionine, N-acetyl cysteine (NAC), a cell permeable glutathione ethyl ester (GEE), and cystathionine reversed the effects of ammonia on T cell proliferation (**Figure 5F and S5F**). A significant increase in ROS and lipid ROS was observed in CD4+ and CD8+ T cells after *in vitro* and *in vivo* treatment with ammonia (**Figure 5G-J**), but not in CRC cell lines (**Figure 5K**). Furthermore, the ferroptotic inhibitor ferrostatin rescued T cell growth inhibition by ammonia (**Figure 5L**). Together, this data indicates that high ammonia levels induce T cell oxidative stress via dysregulation of the transsulfuration pathway.

### Cellular ammonia clearance reduces CRC growth

Ornithine is used clinically in hyperammonemic patients to eliminate ammonia by stimulating both the urea cycle and glutamine synthesis (Davies et al., 2009). Intraperitoneal injection of ornithine at clinically relevant doses to ammonium acetate chow-fed mice reduced serum ammonia and increased T cell proliferation **(Figure 6A and B)**. Moreover, ornithine treatment led to a striking decrease in MC38 and CT26 syngeneic tumor growth (**Figure 6C-D and Figure S6A-B**). The reduction of tumor growth was T cell dependent, as ornithine did not alter growth of MC38 and CT26 tumors implanted in nude mice (**Figure 6E and Figure S6C-E**). Decreased tumor growth was accompanied by significantly reduced Ki67 **(Figure 6F-G and Figure S6F)** and reduced tumor ammonia levels **(Figure 6H and S6G)**. In addition to decreased tumor growth, we also found that ornithine decreased CD44+ tumor stem cells and increased CD4 and CD8 positive T cells in MC38 tumors (**Figure S6H and S6I**). To confirm these findings, an orthogonal approach to reducing systemic ammonia was utilized. Glycerol phenylbutyrate eliminates ammonia via conversion to phenylacetate that binds nitrogen, and conjugates with glutamine to form phenylactylglutamine which is excreted by the kidney (Berry et al., 2014). Glycerol phenylbutyrate treatment significantly reduced MC38 syngeneic tumor size (**Figure S6J and S6K**). To explore the source of the high levels of ammonia, MC38 tumor-bearing mice were treated with lactulose, a microbiota-dependent ammonia reducing agent (Herrmann & Weber, 1987). Lactulose treatment did not significantly reduce tumor size (**Figure S6L**). This suggests that the source of excess ammonia is likely the cancer epithelial cells rather than a dysbiotic microbiota (**Figure S6L**). Ornithine supplementation to the ammonium acetate diet restored T cell function. Ammonia diet reduced T cell secretion of IFNγ, IL-2, and IL-6, but co-treatment with ornithine restored cytokine levels (**Figure S6M**).

**Figure 6:**
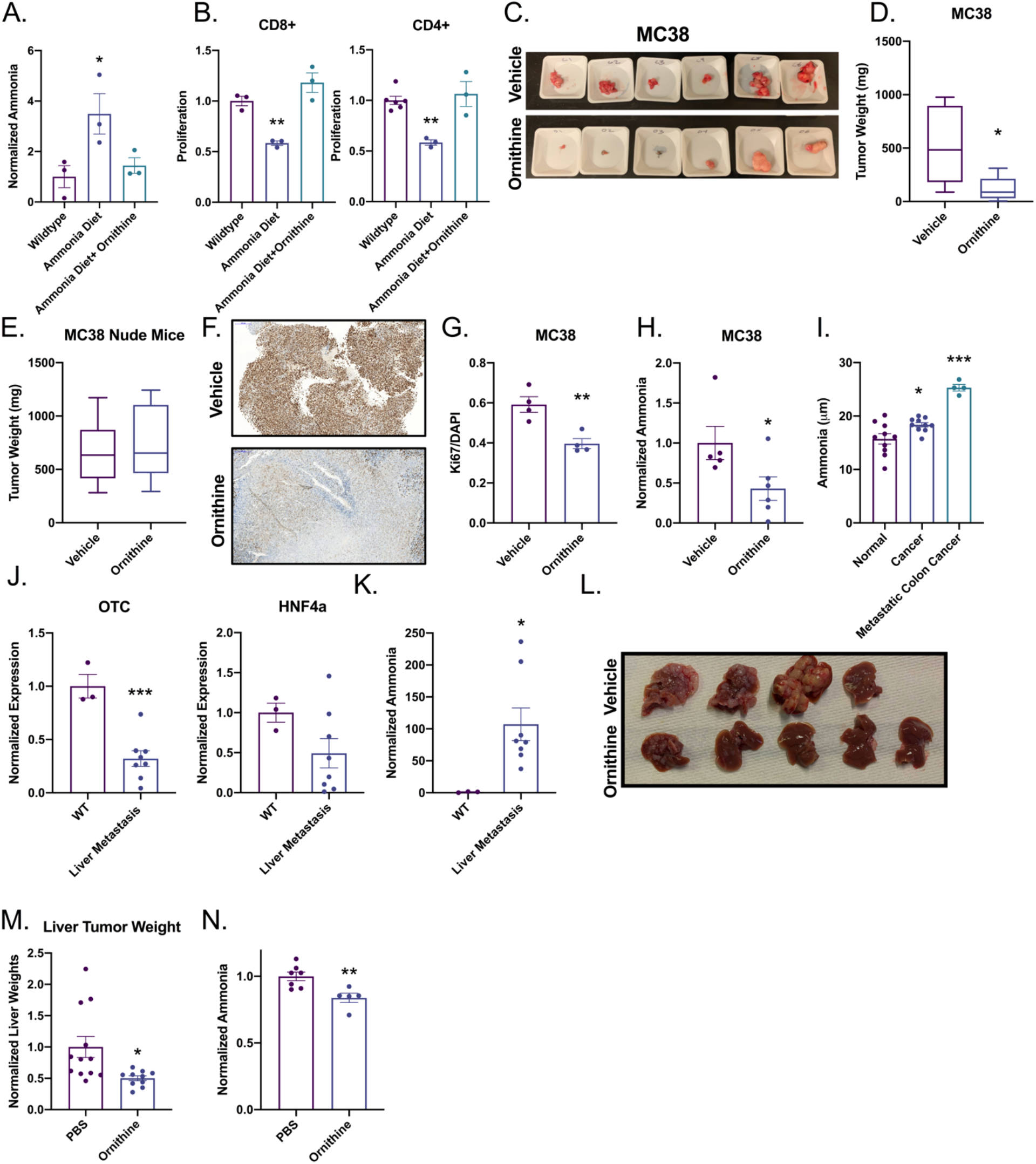
Neutralizing ammonia reduces colorectal cancer tumor growth. A) Serum ammonia or B) T cell proliferation was assessed in wild type mice treated with ammonium acetate diet for 7 days or cotreated with ammonium acetate and daily intraperitoneal injection of 10 mM ornithine (n=3). C,D) Syngeneic tumor study in C57BL/6J or E) athymic nude mice with CRC-derived MC38 cells following ornithine treatment (10 mM ornithine intraperitoneally (n=8-10). Representative IHC F) image and G) quantitation of Ki67 in MC38 tumors treated with ornithine. H) Quantification of ammonia in MC38 tumors treated with ornithine. I) Serum ammonia levels in healthy, CRC, or metastatic CRC patients. J) Gene expression analysis for OTC and HNF4α in normal livers or livers with metastatic lesions K) Ammonia quantification from normal livers or metastatic tumors (n=3). L) Representative images and M) tumor weights of in the liver metastasis model treated with vehicle and intraperitoneal ornithine (n=8-10). N) Ammonia concentration in ornithine treated liver metastasis livers (n=3). *p < 0.05, ** p < 0.01, *** p < 0.001, **** p < 0.0001 unless otherwise indicated. Data is presented as mean +/- the standard error of the mean. All experiments were performed in triplicates at least three times.

To extend these findings to humans, serum ammonia levels were assessed in CRC patients and healthy controls. A significant increase in serum ammonia levels was found in CRC patients compared to healthy controls. Metastatic CRC patient serum ammonia levels were further increased (**Figure 6I**). These results were reflected in the systemic ammonia levels in our murine colorectal cancer model (**Figure S6N**). Having seen a further increase in systemic ammonia levels in metastatic CRC patients, we utilized a previously established syngeneic liver metastasis mouse model (Yu et al., 2021). *Otc* and *Hnf4a* were significantly downregulated in mice with liver metastasis and ammonia levels were increased **(Figure 6J and K.).** Mice treated with ornithine daily had reduced tumor size and maintained more normal liver (**Figure 6L and 6M**). Reduction in tumor size was accompanied by lower ammonia levels, confirming ammonia detoxification (**Figure 6N**).

### High TME ammonia inhibits response to ICB

Ornithine treatment rescues the ammonia driven T cell functional deficits seen in our CRC mouse models. We next evaluated whether ornithine could increase the efficacy of anti-PD-L1 immunotherapy in CRC. *TripleMut* mice were injected with 50 mg/kg tamoxifen, and immediately treated with ornithine or vehicle and co-treated with anti-PD-L1 or isotype control. Anti-PD-L1 or ornithine (p=0.07) alone did not improve mouse survival. However, ornithine and anti-PD-L1 co-treatment combined led to a significant increase in survival (**Figure 7A**). Analysis of ornithine only treated *TripleMut* mice demonstrated that these mice had a significant increase in CD3+ T cell infiltration into the tumors, with less of these T cells being exhausted than the controls (not significant) (**Figure S7A**).

**Figure 7:**
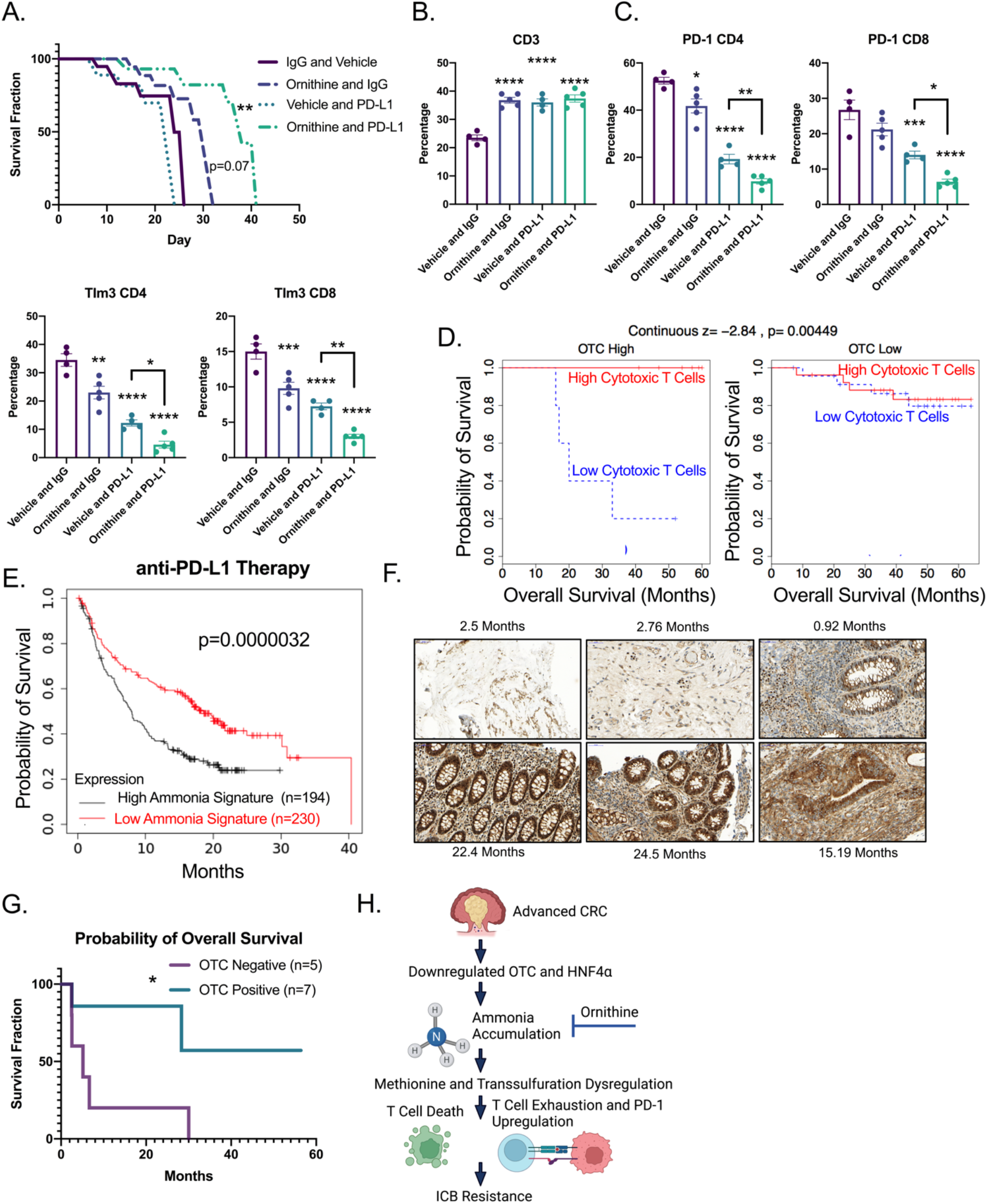
Detoxifying ammonia reactivates immunotherapy in CRC. A) Kaplan-Meier survival curves of *TripleMut* mice injected with 50 mg/kg tamoxifen, and then treated with daily intraperitoneal ornithine, vehicle, isotype control, or anti-PDL1. Significance is compared to vehicle isotype control (n=5-7). B and C) Flow cytometry analysis from *TripleMut* mice treated with ornithine, IgG, or anti-PD-L1 therapy for 7 days. (n=3). D) Human tumor samples were binned into OTC high or OTC low and overall survival was assessed based on cytotoxic T cell scores. (n=62). E) Kaplan-Meier survival curves of PD-L1 treated cancer cohort stratified into high and low ammonia gene signature (n=426). F) Representative OTC staining and G) Kaplan-Meier patient survival based on OTC low and high staining following immunotherapy. Survival significance is calculated using log-rank test. H) Schematic summary of the role of ammonia accumulation in CRC progression. *p < 0.05, ** p < 0.01, *** p < 0.001, **** p < 0.0001 unless otherwise indicated. Data is presented as mean +/- the standard error of the mean. All experiments were performed in triplicates at least three times.

PD-L1 and ornithine co-treatment led to a significant reduction in T cell exhaustion and an increase in overall T cell infiltration (**Figure 7B and 7C and S7B**). We next used our ammonia gene signature to score the data from our RNA-SEQ experiment comparing the *Single, Double*, and *TripleMut* mouse models. A high ammonia score was associated with low CD4 positive T cells in the *TripleMut* (**Figure S7C**). TIDE, an algorithm that correlates gene expression with T cell infiltration, demonstrated that tumors without a strong presence of cytotoxic T lymphocytes, typically have worse survival (Jiang et al., 2018). However, we found that in tumors with low expression of OTC, the presence of cytotoxic T lymphocytes was no longer predictive of survival (**Figure 7D**). In addition, we found that a gene signature score associated with high ammonia levels predicted worse survival following ICB treatment in a large cohort of combined cancer types (**Figure 7E**). We assessed a small institutional cohort of MSI-high CRC patients treated with immunotherapy. Patients with low OTC staining (associated with high ammonia levels) by IHC had worse survival after PD-1 treatment (**Figure 7F and 7G**). OTC levels were not associated with differential response to traditional chemotherapy (**Figure S7D**). We hypothesize that in advanced CRC, OTC and HNF4α are downregulated which leads to ammonia accumulation, transsulfuration dysregulation in T cells, increased T cell exhaustion and ICB resistance (**Figure 7H**).

## DISCUSSION

Altered cellular metabolism is a hallmark of cancers and impacts the tumor microenvironment, immune landscape, and therapy response. Moreover, metabolic reprogramming in tumors generates excess byproducts that must be counteracted by a heterocellular metabolic exchange in the tumor microenvironment. Here, using a highly metastatic model of CRC (*TripleMut*), we show that there is a robust downregulation of T cell activation and function compared to more benign less progressive CRC mouse models (*DoubleMut* or *SingleMut* mice). These data are consistent with other models of CRC demonstrating a robust immunosuppressive tumor microenvironment (W. Liao et al., 2019). Through a multi-omic analyses integrating transcriptomic, CyTOF and unbiased metabolomics data, we identified a novel connection between ammonia metabolism and T cell function. The activity of HNF4α and its downstream target genes involved in ammonia clearance were robustly inhibited in CRC, leading to a buildup of extracellular ammonia. Consistent with the mouse model data, *HNF4a* and the expression of ammonia clearance genes were downregulated and serum ammonia was increased in CRC patients. High extracellular ammonia decreased T cell anti-tumor function, while not altering viability or function of other immune cells. Using several orthogonal approaches, neutralizing ammonia levels decreased tumor growth in a T cell dependent manner, whereas increasing tumor ammonia led to attenuation of T cell function and increased progression in the *SingleMut* mice.

A nutrient limited microenvironment alters T cell proliferation and function. Insufficient oxygen and waste removal occurs as tumors grow. Hypoxia inhibits CD4+ T cell function and promotes regulatory T cell activity (Westendorf et al., 2017) while accumulation of lactate suppresses cytotoxic T lymphocytes (Wang et al., 2021). Previous work has shown that ammonia accumulates in hepatocellular carcinoma (Bai et al., 2021). Urea cycle dysregulation is a common feature of tumors, and ammonia is a central nitrogen source to support cancer growth (Spinelli, Yoon, et al., 2017). However, little work has been performed on how altered ammonia metabolism in cancers affect stromal cells (Lee et al., 2018). Our data reveals that dysregulation of urea cycle enzymes is correlated with increased extracellular ammonia, which is a major mechanism that leads to an immunosuppressive tumor microenvironment in CRC. The tumor microenvironment is an important driver of immunosuppression and response to ICBs (Bader et al., 2020).

Recent work suggests a more complicated metabolic interaction where nutrients are selectively partitioned to immune and cancer cells based on uptake mechanisms (Reinfeld et al., 2021). However, decreases in glutamine, lipids, and amino acids in the tumor microenvironment limit T cell growth, viability and/or function (Bian et al., 2020; Chang & Pearce, 2016). Our metabolomics data from ammonia treated T cells implicated altered methionine metabolism. T cells proliferate rapidly and have a high metabolic requirement for methionine. Recent work has shown that methionine is necessary to drive T cell differentiation and proliferation (Sinclair et al., 2019). Methionine can be converted to S-adenosylmethionine which is a common methyl donor. Tumors disrupt methionine metabolism and reduce S-adenosylmethionine in CD8+ T cells, which leads to impaired T cell immunity (Bian et al., 2020). Our work highlights the requirement of the transulfuration pathway for the conversion of methionine to cysteine, which is the limiting substrate for glutathione synthesis. Flux analysis revealed that ammonia decreased transsulfuration flux and lower glutathione levels, that were associated with increased ROS in T cells. The robust ammonia-induced expression of *Bhmt*, would inhibit the transsulfuration pathway by siphoning off homocysteine for methionine synthesis. Although the mechanism by which *Bhmt* is induced is unclear, we speculate that this may be a strategy for conserving the methionine pool in T cells. However, a chronic increase in BHMT is predicted to be detrimental, leading to a lower glutathione pool size, higher ROS accumulation and induction of ferroptosis.

Although some work has shown that an immunoscore for T cell infiltration predicts survival, initial CRC approaches to ICB have been very disappointing (Ganesh et al., 2019). The majority of CRC is microsatellite stable, but 15% of cancer has high microsatellite instability (MSI) (Gatalica et al., 2016). However, in MSI high CRC, the objective response after PD-1 inhibitor is 40% compared to 0% in non MSI high tumors (Brahmer et al., 2010). Most CRCs are not MSI high and there is significant heterogeneity in immune infiltration (Picard et al., 2020). Our work demonstrates that clearance of ammonia decreases tumor growth and potentiates ICB response in CRC models that are refractory to immunotherapy. High OTC expression in patient samples is associated with ammonia clearance, and an improved response to immunotherapy. The data, including work done in T cell deficient models, points to an ammonia/T cell-axis that is central in CRC growth and the immunotherapy response. Reactivating ammonia clearance may be a viable therapeutic option to increase response to ICB in non MSI high CRC. However, more functional studies are needed to understand how ammonia in the tumor microenvironment may alter the numbers and/or function of other stromal cells.

Ammonia metabolism and accumulation is unique in colon cancer compared to other cancers due to two major sources of ammonia: cell autonomous metabolism and the microbiota. Many commensal microbes including *Clostridia, Enterobacteria*, and *Bacillus spp* produce large quantities of ammonia (Vince and Burridge, 1980) predominantly by amino acid deamination and urea hydrolysis by urease (W. Zhang et al., 2021). Host cells generate ammonia via the conversion of L-glutamine to L-glutamate catalyzed by glutaminase (Cooper & Jeitner, 2016). Ammonia is largely eliminated through the urea cycle. Ammonia accumulates rapidly in tumors due to poor vascularization and increased glutamine metabolism. Moreover, *Kras* mutations are known to drive glutamine metabolism, which is in line with *TripleMut* mice having the highest level of tumor ammonia and our most progressive and immunosuppressive model. Interestingly, activation of the *Kras* mutation by itself did not lead to increased ammonia levels or dysregulation of HNF4α activity. We hypothesize that the metabolic changes leading to high ammonia result from the combinatorial effects of the *Apc, Kras*, and p53 mutations*Tp53*, which will be tested using the appropriate mouse models in the future.

Our work suggests the tumor protection derived from ammonia detoxification is T cell dependent. The temporal studies done *in vivo* suggest that initially, increased ammonia levels led to T cell exhaustion and increased expression of PD-1 and CTLA-4 on T cells. As the tumors progress, ammonia levels accumulate leading to a reduction in T cell numbers. The same phenomenon is seen *in vitro*, as markers of T cell exhaustion are upregulated early and prior to changes in cell viability. Higher levels of ammonia *in vitro* potently reduce T cell growth and lead to cell death. That said, more studies will be helpful in informing the time course of ammonia accumulation on T cell numbers and function *in vivo*. This is difficult to evaluate in late stage *TripleMut* mice due to the rapid progression of disease, and variability between mice. Detoxifying ammonia via ornithine treatment increased T cell numbers in a syngeneic CRC model. In the *TripleMut* model, which we consider to be more progressive and more immunosuppressive, we see that ornithine alone reduces T cell exhaustion markers and increases T cell numbers. In this advanced model, PD-L1 therapy alone is largely ineffective. Combined ornithine and PD-L1 therapy combined leads to a significant increase in total T cells and T cell activation, and a further decrease in markers of T cell exhaustion. The combination of ornithine and anti-PD-L1 immunotherapy also increased survival by 72%, indicating that the two-pronged approach to reactivate T cells is effective. Low levels of ammonia signature are associated with improved survival after anti-PD-L1 therapy in a large cohort of many cancer types. In addition, high OTC expression is associated with improved survival after ICB treatment in patients with MSI-high CRC.

In sum, our results reveal dysregulation of the urea cycle in CRC, which leads to extracellular ammonia accumulation. Ammonia preferentially suppresses T cell growth and induces exhaustion and immunosuppressive markers. Ammonia detoxification is therefore a potential novel therapeutic option, particularly for metastatic and non-MSI high CRC.

## METHODS

### Experimental Model and Subject Details

#### Mice

All mice used in this paper are in a predominant C57Bl/6J background. Males and females are equally represented and littermates were randomly mixed in all experimental conditions. The mice were housed in a temperature controlled, specific pathogen free environment, with a 12 hour light/dark cycle. They were fed ad libitum with standard chow diet. Studied mice were between 4-6 weeks old. Mouse lines used were CDX2-ER^T2^Cre; *Apc^fl/fl^* mice; ROSA *KRAS*^G12D^ *CDX2*-ER^T2^Cre; *Apc^fl/fl^* mice, CDX2-ER^T2^Cre; *Apc^fl/fl^*; *Tp53^fl/fl^*, and *Apc^fl/fl^;Tp53^fl/fl^*; Kras^LSLG12D^ mice. All animal studies were carried out in accordance with Association for Assessment and Accreditation of Laboratory Animal Care International guidelines and approved by the University Committee on the Use and Care of Animals at the University of Michigan.

#### Human Subjects

The study was approved by the institutional review boards at the University of Michigan (IRBMED, protocol number HUM00085066 and HUM00064405) and the Department of Health and Human Services, Food and Drug Administration (Research Involving Human Subjects Committee/RIHSC, protocol number14-029D). All protocol and applicable local regulatory requirements were meant. All adult male and female subjects provided written informed consent and review of their medical history.

#### Human Survival Analysis

Human survival analysis was determined using the online tool KMplot (Lánczky & Győrffy, 2021). This is a web based, registration-free survival analysis tool that can perform univariate and multivariate survival analysis. For both general survival and immunotherapy survival, we used our ammonia gene signature as an input. Significance is computed using the Cox-Mantel log rank test.

#### Cell Lines

Human intestinal cell lines HCT116, RKO, SW480, and DLD1 were used for most experiments. Other cell lines used are MC38 and CT26. Cell lines have been STR-authenticated. All cells were maintained in complete DMEM medium (supplemented with 10% fetal bovine serum and 1% antibiotic/antimycotic agent) at 37°C in 5% CO_2_ and 21% O_2_.

### Method Details

#### Growth Assay

Adherent cell growth assays were performed using live cell imaging using the Cytation 5 Imaging Multi-Mode reader. Cells were plated down, treated 24 h later with indicated treatments, and immediately imaged and analyzed for cell number. Images were then taken every 24 h. Cytation software was used to quantify cell number.

#### Real time Quantitative PCR

One μg of total RNA, extracted using Trizol reagent from mouse tissues (intestinal epithelial scrapes), human intestine, primary T cells and cell lines, was reverse transcribed to cDNA using SuperScriptTM III First-Strand Synthesis System (Invitrogen). Real time PCR reactions were set up in three technical replicates for each sample. cDNA gene specific primers, SYBR green master mix was combined, and then run in QuantStudio 5 Real-Time PCR System (Applied BioSystems). The fold-change of the genes were calculated using the ΔΔCt method using *Actb* as the housekeeping gene mRNA. Primers are listed in **Table S4**.

#### Metabolomics

For indicated mouse model, mouse was induced with Tamoxifen, and colon scrapes were collected day 10 after induction. Colon scrapes were snap-frozen in liquid nitrogen and then normalized by mass. Metabolites were extracted by adding dry-ice cold 80% methanol for 10 minutes before bead homogenization. Samples were clarified via high speed centrifugation and supernatant extracted. Metabolite extracts were then lyophilized using a SpeedVac concentrator and resuspended in 50:50 methanol/water mixture for LC-MS analysis. For T-cell samples, cells were treated at 10 million cells per .5 ml of complete RPMi. Samples were then washed once with ice cold PBS, then incubated in dry-ice cold 80% methanol on dry ice for 10 minutes before homogenization. Samples were normalized by conducting BCA protein concentration on a technical replicate and volume normalizing across samples to achieve even concentrations. Metabolite extracts were then lyophilized using a SpeedVac concentrator and resuspended in 50:50 methanol/water mixture for LC-MS analysis.

Data was collected in both negative and positive ion modes. For negative mode, previously published parameters were used (Bennett et al., 2008; T. U. Chae et al., 2015; Singhal et al., 2021). For positive mode, an Agilent Technologies Triple Quad 6470 LC/MS system consisting of a 1290 Infinity II LC Flexible Pump (Quaternary Pump), 1290 Infinity II Multisampler, 1290 Infinity II Multicolumn Thermostat with 6 port valve and 6470 triple quad mass spectrometer was used. Agilent Masshunter Workstation Software LC/MS Data Acquisition for 6400 Series Triple Quadrupole MS with Version B.08.02 is used for calibration, compound optimization and sample data acquisition. A Waters Acquity UPLC HSS T3 1.8 μm VanGuard Pre-Column 2.1 x 5 mm column and a Waters UPLC BEH TSS C18 column (2.1 x 100mm, 1.7μm) was used with mobile phase A) consisting 0.1% formic acid in water; mobile phase (B) consisting of 0.1% formic acid in acetonitrile. The solvent gradient was: mobile phase (B) held at 0.1 % for 3 min, increased to 6 % at 12 min, 15 % at 15 min, 99% at 17 min and held for 2 min before going to initial condition and held for 5 min. The column was held at 40 °C and 3 μl of sample was injected into the LC-MS with a flow rate of 0.2 ml/min. Calibration of 6470 QqQ MS was achieved through Agilent ESI-Low Concentration Tuning Mix.

QqQ data for both ion modes were preprocessed with Agilent MassHunter Workstation Quantitative Analysis Software (B0700). Metabolite chromatogram peaks were manually inspected. Each metabolite abundance level was median normalized across the sample population. Statistical significance was determined by a one-way ANOVA with a significance threshold of 0.05. Metaboanalyst was used for pathway enrichment analysis.

##### [^35^S]-Methionine flux into GSH

T cells (15 million cells in 0.5 mL of RPMI media) were cultured in a 12-well plate. L-[^35^S]-methionine (1175 Ci/mmol, NEG009A500UC, Perkin Elmer), was added (5 μCi/ml) and incubated for 24 h. Then, the T cell suspension was transferred to 1.5 ml tubes and centrifuged at 10,000 x *g* for 3 min. The supernatant was discarded, and the cell pellet was suspended in 150 μl of 1X PBS. One aliquot of the suspension (100 μl) was mixed with an equal volume of metaphosphoric acid solution (16.8 mg/ml) with EDTA and 150 mM NaCl while a second aliquot (30 μl) was mixed with equal volume of RIPA lysis buffer (R0278, Sigma) with protease inhibitor cocktail for mammalian tissue extract (P8340, Sigma) for protein quantitation using the Bradford reagent. The RIPA lysis buffer and protease inhibitor is mixed in the ratio of 100:1. After vortexing for 5-10 sec, the samples were frozen on dry ice and stored at −20 °C. For HPLC analysis, samples were thawed on ice, centrifuged at 12,000 x *g* for 5 min and mixed with 15 μl of monoiodoacetic acid. The pH was adjusted to 7-8 (monitored using pH paper strips) and incubated for 1 h in the dark at room temperature. Then, the samples were mixed with an equal volume of Sangers reagent (15 μl/ml in absolute ethanol, 200 proof, Decon Laboratories), incubated at room temperature for 4 h and separated by HPLC using a Bondapak column (Agilent) as described previously(Mosharov et al., 2000).

#### Ammonium Acetate Diet

Diet was made from pelleted chow mixed with 25% powdered ammonium acetate and water. It was mixed using a KitchenAid food mixer and dried in a dehydrator.

#### IL 17 Bone Marrow Transplant

Eight-week-old mice from the indicated genotype were lethally irradiated twice on the same day. Bone marrow was isolated from the indicated genotype from a separate donor mouse, and injected into the tail vein on the same day. Mice were monitored for diarrhea and weight loss daily.

#### T Cell Isolation

CD4 and CD8 T cells were isolated from the spleen. Briefly, the spleen was removed from a sacrificed mouse, pushed through a sterile 70 micron filter using the plunger of a syringe, and then washed in PBS. Cells were incubated with the antibody and beads from the MojoSort CD3, CD4 and CD8 T cell isolation kit, and then negative selection was achieved using a magnet. Cells were counted and plated in RPMI with FBS. T cell proliferation was performed under plate-bound CD3 and CD28 stimulation. Cells were incubated for 45 minutes with MTT solution (5X concentrate stock: 5 mg/ml, in 1XPBS, pH 7.4) or 15 minutes woth CFSE (5 μM). Absorbance was read at 570nm for MTT and CFSE was assessed by the Z2 flow cytometer.

#### CyToF

The indicated mouse line was induced with tamoxifen and killed on day 10 after induction. The colons were incubated with 10 mM EDTA and the epithelial cells were discarded. The colons were next incubated with 1 mg/ml of Type IV collagenase for 40 minutes, and then immune cells were isolated using a Percoll gradient. Cells were stained with indicated CyToF antibodies (Table 1), and a DNA intercalator after permeabilization with paraformaldehyde. Samples were acquired on the Helios (Fluidigm) and analyzed in CytoBank.

#### Ammonia Quantification

Cells or tissues were deproteinated using a water-methanol-chloroform gradient. Lysates were then centrifuged rapidly to clear debris. Ammonia was then quantified using the L-Glutamate Dehydrogenase ammonia kit (Sigma). Ammonia was converted to indophenol using the Berthelot reaction, and then quantified using mass spectrometry as previously described (Spinelli, Kelley, et al., 2017).

#### C11-BODIPY lipid ROS measurement

Wildtype splenic T cells or CRC cells were incubated for 24 h with the indicated concentration of ammonia. Cells were harvested in HBSS, washed, and stained with 5 μM C11-BODIPY (ThermoFisher) at 37 °C for 30 minutes. Fluorescent intensity was measured on the FITC channel on the Z2 flow cytometer. A minimum of 20,000 cells were analyzed per condition, data was analyzed using FlowJo software (Tree Star). Values are MFI.

#### ROS Detection Assay

ROS detection assay was performed as previously described(Bell et al., 2021). Briefly, cell permeable free radical sensor carboxy-H2DCFDA (Invitrogen) was used. Cells were treated with indicated ammonia concentration for 24 h, then incubated with 10 μM carboxy-H2DCFDA in PBS at 37 °C for 45 min. Cells were then washed and resuspended in PBS. Mean fluorescent intensity was obtained from the Cytation 5 Imaging Multimode Reader.

#### Histology

Tumor tissues were rolled and fixed in formalin for 24 h, then embedded in paraffin. Sections of 5 μm were stained for H&E and mounted with Permount Mounting Medium (Thermo Fisher Scientific). For immunohistochemistry, paraffin tissue sections underwent antigen retrieval, blocking in 5% goat serum in PBS, and probed with Ki67 antibody (Cell Signaling, 1:250), OTC antibody (ThermoFisher 1:200), or CD8 antibody (BD Bioscience 1:200). Sections were washed twice with PBST and incubated with HRP conjugated anti-rabbit IgG (1:500, catalog 7074S, Cell Signaling Technology) for 1 h. Sections were then washed with PBST and stained with DAB substrate. After the color change, the reaction was stopped with distilled water and dehydration was completed before mounting with Permount Mounting Medium. For immunofluorescence OPAL staining, samples were initially dehydrated, then stained with the indicated fluorescent OPAL antibody, then analyzed by multispectral imaging and analyzed for spatial and morphological context. Histological scoring of dysplasia was done by a blinded pathologist.

#### Nessler’s Reagent Staining

Nessler’s reagent staining was performed as described in (Gutiérrez-de-Juan et al., 2017)

#### Immune Cell Identification

Colons were removed, washed thoroughly in PBS, and then disassociated with 10 mM EDTA for 45 minutes. The remaining colon after this treatment was incubated with 1 mg/mL collagenase Type IV for 45 minutes. Immune compartment was obtained through a Percoll gradient, cells were stained using indicated antibody, and then acquired using flow cytometry.

#### RNA-SEQ analysis

Indicated mouse line was induced with tamoxifen and tissues were collected on day 10 after injection. Colons were washed gently in PBS, scraped with a glass slide, and then the scraped tissue was flash frozen with liquid nitrogen. RNA was isolated from tissue as described above. Quality of fastq files was assessed using FastQC v0.11.8 and MultiQC v1.7(Ewels et al., 2016). Reads were then aligned to GENCODE’s GRCm38.vM24 assembly using STAR v2.6.1a_08-27(Dobin et al., 2013; Frankish et al., 2018). Aligned reads were counted using featureCounts v1.6.3(Y. Liao et al., 2014). PCA-based clustering was used to identify outliers and then differential expression was performed using DESeq2 v1.30.1(Ren & Kuan, 2020). Genes with an adjusted p-value < 0.05 were considered differentially expressed. All analysis was carried out on the University of Michigan Great Lakes HPC cluster.

### Survival analysis

The TCGA data was DESeq normalized and the mean expression of the genes included in the signature was used in the survival analysis employing overall survival data. The differential survival was assessed by a Cox proportional hazards regression and Kaplan-Meier plots were drawn to visualize the survival difference. The plots also include hazards rates and 95% confidence intervals.

#### Syngeneic and Xenograft Studies

Wildtype or athymic nude mice of both sexes were inoculated with 2 million MC38 or CT26 cells. Cells were implanted into lower flanks, and treatment began at day 10 after palpable tumors. Tumor size was measured with digital calipers. At the endpoint roughly 30 days or until tumors were 2 cubic centimeters, mice were killed and tumors excised. Tumor volume and weight was measured and tissues were prepared for histology, IHC, and flow cytometry.

#### Liver Metastasis Model

Liver metastasis model was performed as previously described(Yu et al., 2021). Briefly, mice were anesthetized using isoflurane, a small dorsal right side incision was made and the splenic artery was located. Cells resuspended in PBS were injected into splenic artery which was sutured shut and splenectomy was performed. The mouse fascia and skin was then closed.

### Quantification and Statistical Analysis

*In vitro* experiments were validated in multiple cell lines. Each condition of each cell line experiment was performed in technical replicates and repeated three times to ensure reproducibility. Figures show a representative biological replicate unless otherwise indicated. Blinding was performed whenever appropriate. Sample description and identification was unavailable to the core personnel during data collection and analysis. Statistical details of all experiments can be found in the figure legends. The sample numbers are mentioned in each figure legend and denote biological replicates. Statistical details are reported in the figure legends. Results are expressed as the mean +/- standard error of the mean for all figures unless otherwise noted. Significance between 2 groups was tested using a 2 tailed unpaired t test. Significance among multiple groups was tested using a one-way ANOVA. GraphPad Prism 7.0 was used for the statistical analysis. Statistical significance is described in the figure legends as: * p < 0.05, ** p < 0.01, *** p < 0.001, **** p < 0.0001.

## Acknowledgements

This work was funded by NIH grants: R01CA148828, R01CA245546, and R01DK095201 (Y.M.S); R37CA237421, R01CA248160, and R01CA244931 (C.A.L); UMCCC Core Grant P30CA046592 (Y.M.S and C.A.L); and R35GM130183 (R.B). H.N.B was supported by T32 training grant (GM008322) and F30CA257292. R.K was supported by a postdoctoral fellowship from the American Heart Association (826245) and NIH grant F30CA257292. S.A.K was supported by an NIH F31 fellowship (F31CA247457). S.L.M was supported by CMB Graduate Program T32GM007315. Metabolomics studies were supported by NIH grant DK097153, and the Charles Woodson Research Fund. R.S was supported by an American Physiological Society postdoctoral fellowship. S.S was supported by the Crohn’s and Colitis Foundation Research fellowship award (623914) and the American Heart Association postdoctoral fellowship (19POST34380588). B.G was supported by the 2020-1.1.6-JÖVŐ-2021-00013 and 2020-4.1.1.-TKP2020 grants.

## Conflict of Interest

C.A.L. has received consulting fees from Astellas Pharmaceuticals, Odyssey Therapeutics, and T-Knife Therapeutics, and is an inventor on patents pertaining to Kras regulated metabolic pathways, redox control pathways in pancreatic cancer, and targeting the GOT1-pathway as a therapeutic approach (US Patent No: 2015126580-A1, 05/07/2015; US Patent No: 20190136238, 05/09/2019; International Patent No: WO2013177426-A2, 04/23/2015).

## Notes

The authors declare no potential conflicts of interest.

### Competing Interest Statement

The authors have declared no competing interest.

