## Supplemental Figures for "Microenvironmental Ammonia Enhances T cell Exhaustion in Colorectal Cancer"

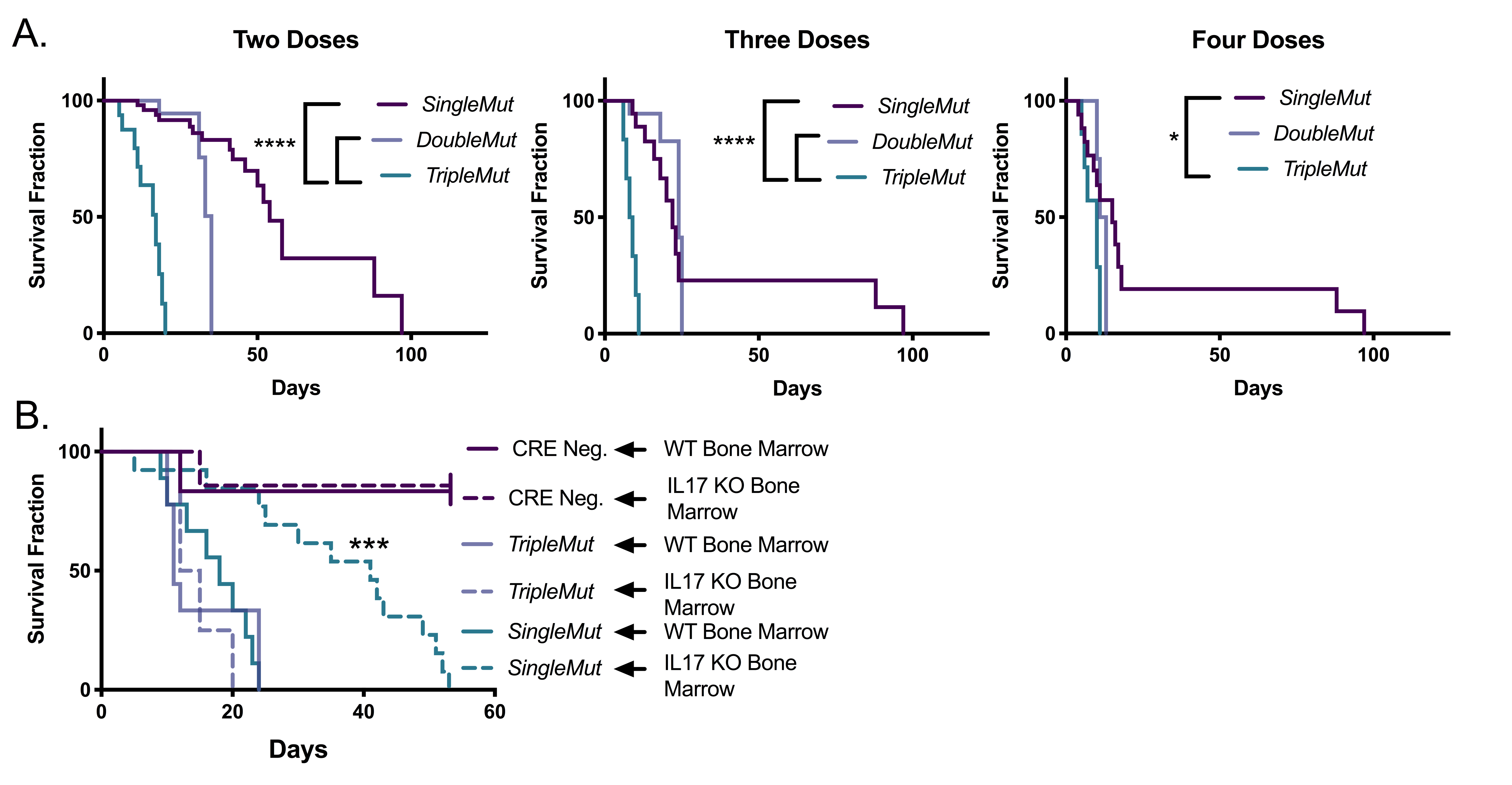
**Figure S1: Dose dependent survival in *SingleMut*, *DoubleMut*, and *TripleMut* mouse model.** A) Mice were induced with the indicated doses of 50 mg/kg tamoxifen and then followed for survival (n=8-10). B) Irradiated mice were i.v. injected with the indicated bone marrow, then followed for survival. (n=4-6). Survival significance is calculated using log-rank test. *p < 0.05, ^∗∗^ p < 0.01, ^∗∗∗^ p < 0.001, ^∗∗∗∗^ p < 0.0001 unless otherwise indicated. Data is presented as mean +/- the standard error of the mean.

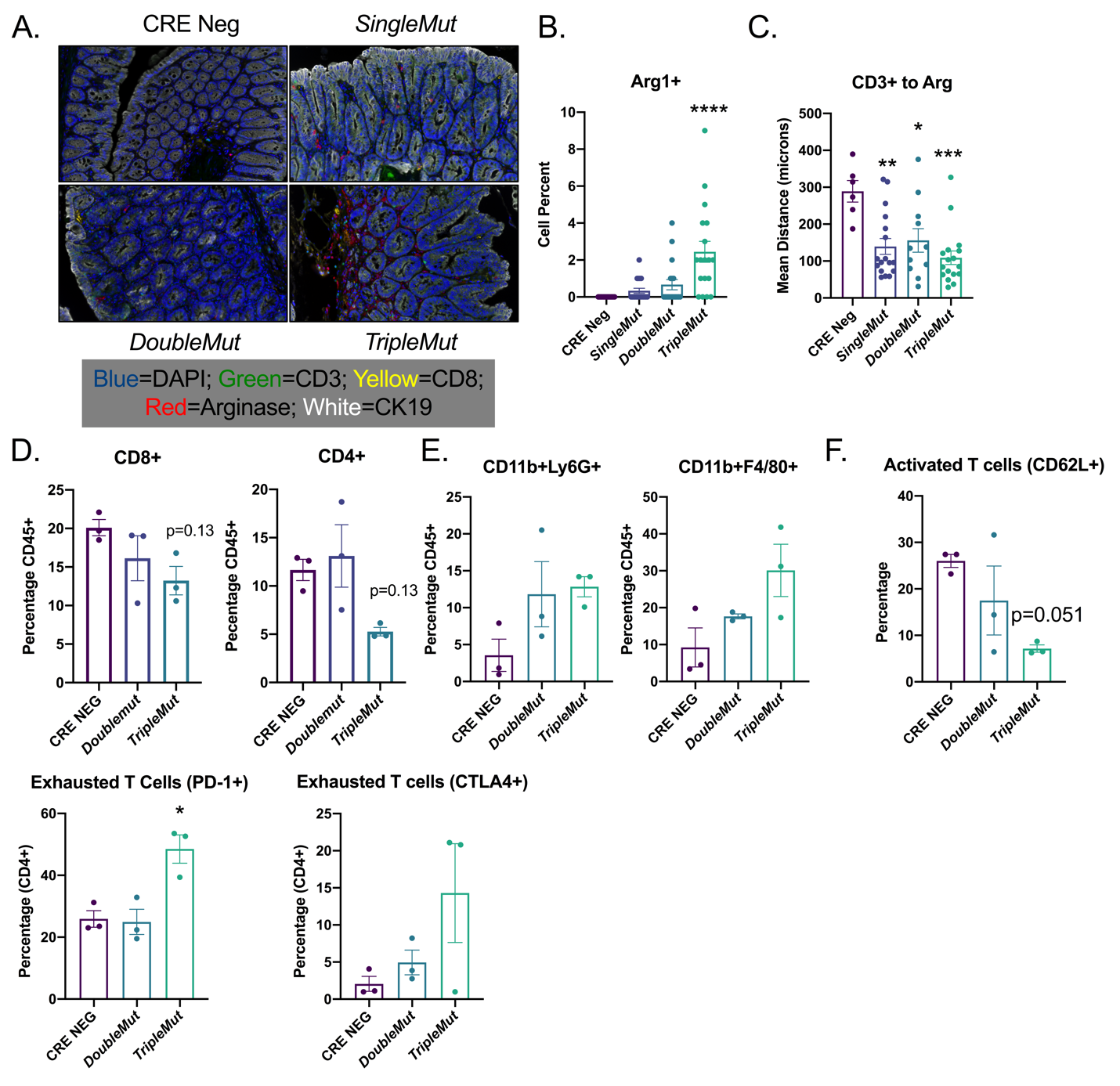

**Figure S2: *TripleMut* mice have altered tumor microenvironment.** A) Representative OPAL multiplex IHC in mouse models of colon cancer induced with 100 mg/kg tamoxifen. B) Quantitation of Arg1+ cells (n=6-8). C) Average distance between cell populations (n=6-8). D, E, and F) Selected populations from CyToF experiment (n=3). G) Flow on end-stage *DoubleMut* and *TripleMut* mice (n=4-6). *p < 0.05, ^∗∗^ p < 0.01, ^∗∗∗^ p < 0.001, ^∗∗∗∗^ p < 0.0001. Data is presented as mean +/- the standard error of the mean.

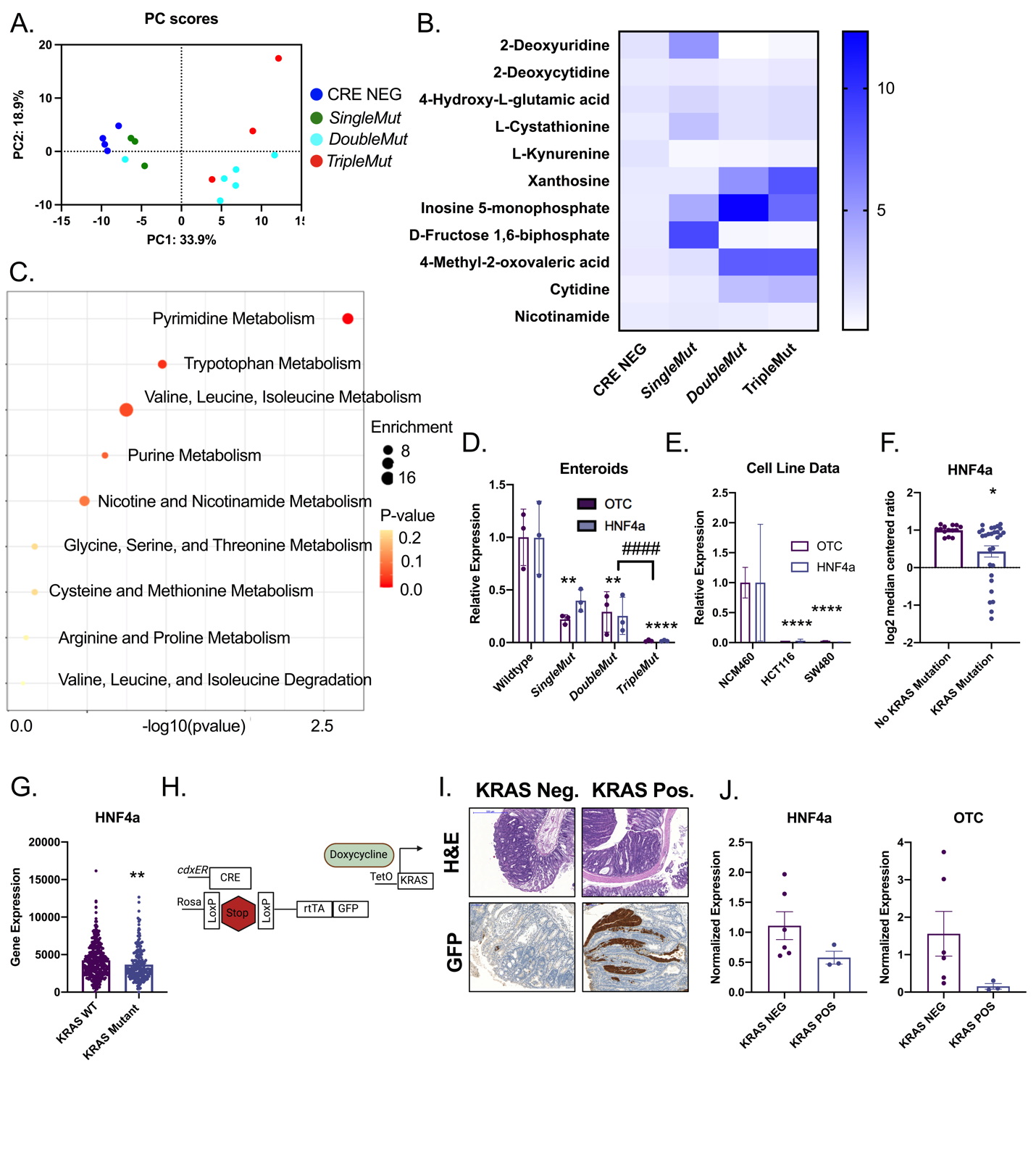

**Figure S3: OTC and HNF4**α **are dysregulated in *KRAS* mutant colorectal cancer.** Indicated mice were induced with 3 injections of 100 mg/kg tamoxifen. Colon scrapes were isolated for metabolomics and A) PCA analysis, B) significant metabolite changes , and C) pathway enrichment analysis (n=3-6). D) Expression of OTC and HNF4α in enteroids from indicated mouse models. E) Expression of OTC and HNF4α in colorectal and normal colon cell lines. Expression of HNF4α in F) cell lines or G) CRC patient samples stratified by *Kras* mutational status. H) Schematic depicting the *iKras* mouse model. I) Representative H&E and GFP immunohistochemistry and J) gene expression for HNF4α and OTC from *Kras* mutant and *Kras* wild type mice. *p < 0.05, ^∗∗^ p < 0.01, ^∗∗∗^ p < 0.001, ^∗∗∗∗^ p < 0.0001 unless otherwise indicated. Data is presented as mean +/- the standard error of the mean.

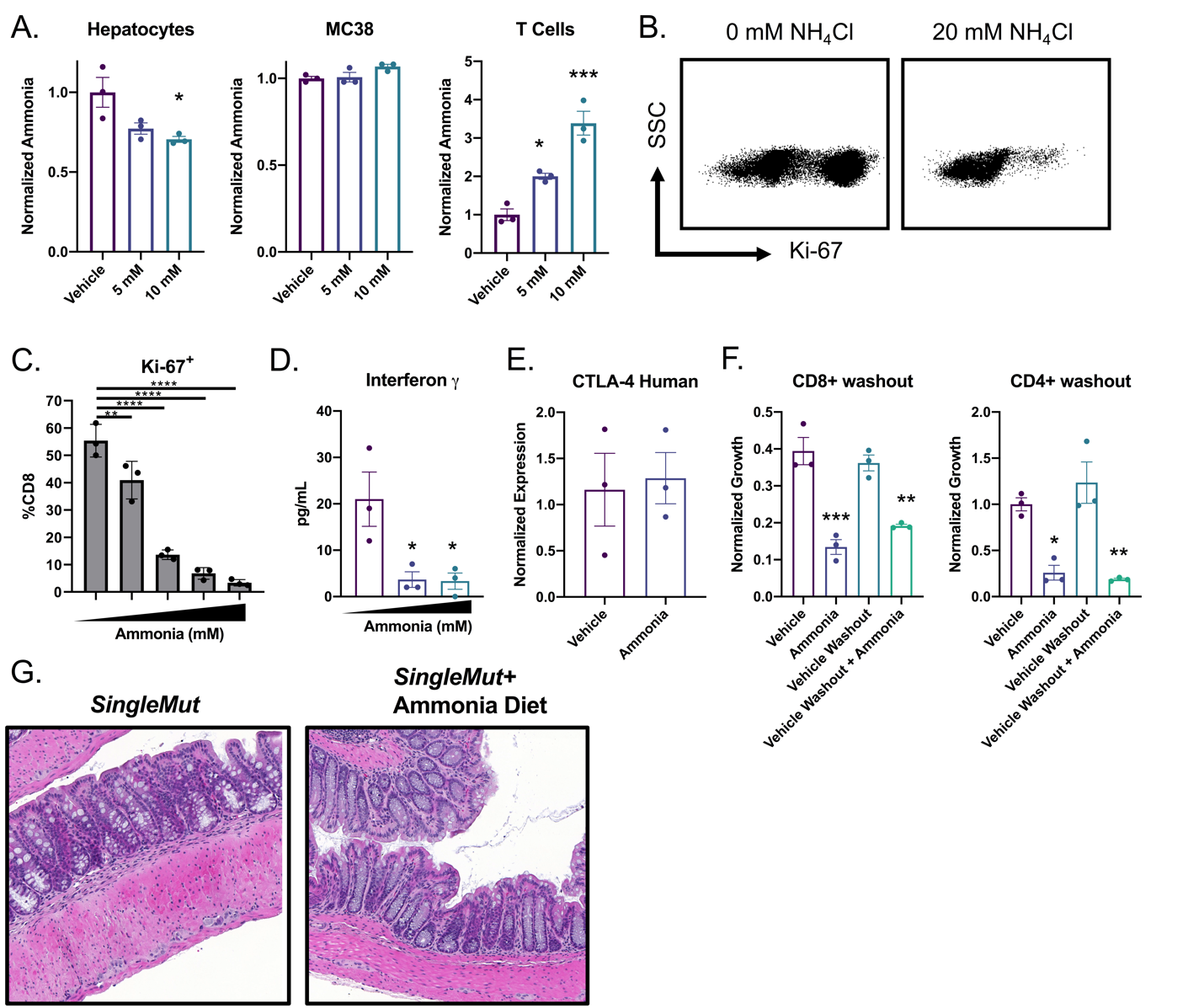

**Figure S4: Ammonia induces a durable reduction in proliferation.** A) Indicated cells were treated for 24 hours with ammonia, then washed with PBS, lysed, deproteinated, and an ammonia assay was performed. B and C) Representative plot and quantification of Ki67 expression in ammonia-treated CD8+ T cells. D) ELISA of supernatant of ammonia treated T cells after 12 h ammonia treatment. E) Gene expression analysis in human T cells treated with ammonia for 72 h. F) T cells were plated with ammonia for 24 h, then the ammonia was washed off and T cells were replated in fresh media with no ammonia then assessed for proliferation. G) Representative H and E of *SingleMut* on ammonium acetate diet. *p < 0.05, ^∗∗^ p < 0.01, ^∗∗∗^ p < 0.001, ^∗∗∗∗^ p < 0.0001 unless otherwise indicated. Data is presented as mean +/- the standard error of the mean. All experiments were performed in triplicates at least three times.

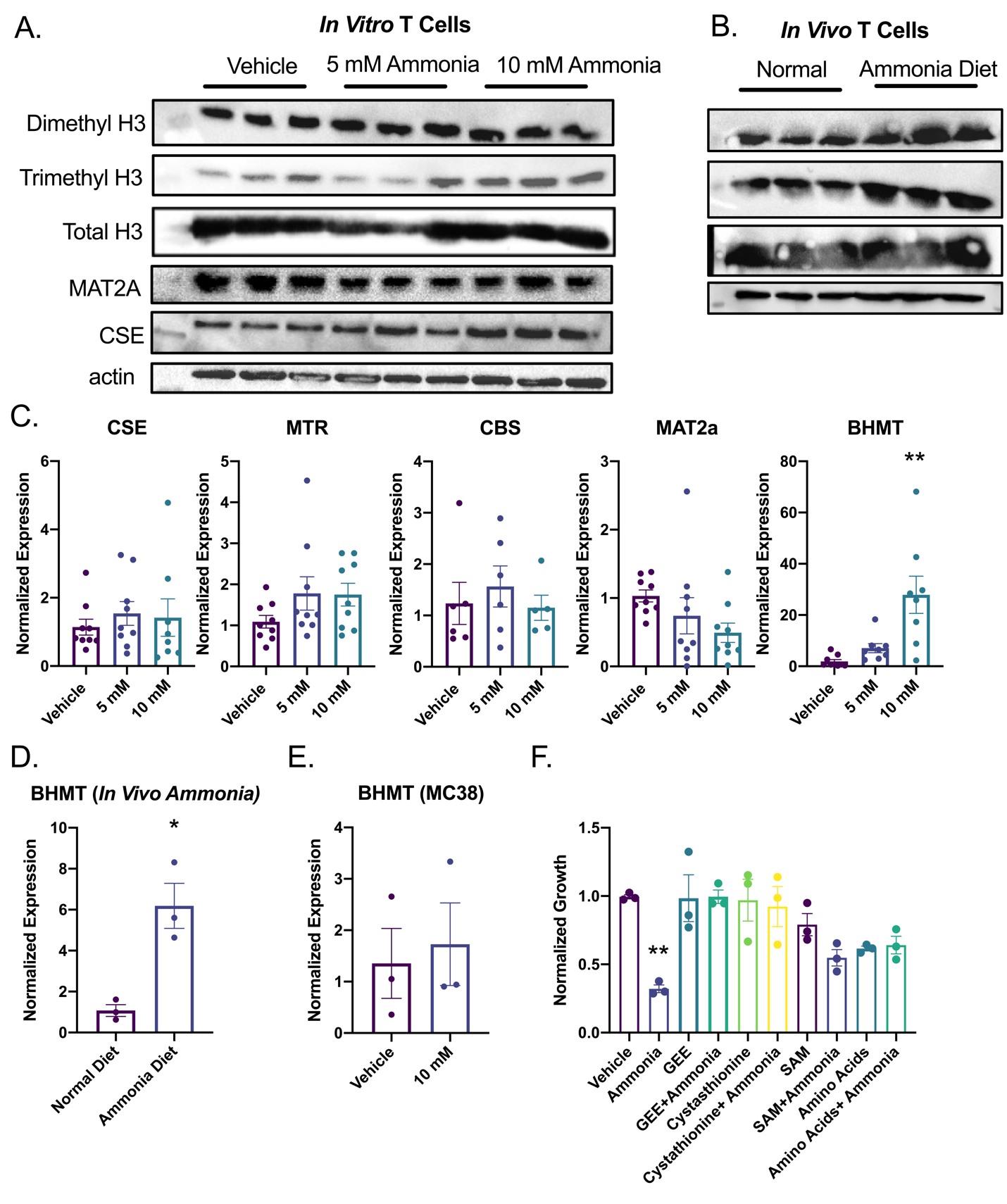

**Figure S5: Ammonia upregulates BHMT expression in T cells.** A and B) Western on *in vivo* and *in vitro* ammonia treated T cells for indicated antibody (n=3). T cells were treated with indicated ammonia concentration C) for 24 h or D) ammonia acetate diet for 7 days and gene expression was assessed. E) MC38 cells were treated with 10 mM ammonia for 24 h, then analyzed for gene expression. F) T cells were treated with indicated compound for 24 h (GEE, 1 mM; SAM, 50 μm; cystathionine, 500 μm; amino acids, 200 μm) then incubated with 10 mM ammonia for 24 h. *p < 0.05, ^∗∗^ p < 0.01, ^∗∗∗^ p < 0.001, ^∗∗∗∗^ p < 0.0001 unless otherwise indicated. Data is presented as mean +/- the standard error of the mean. All cell experiments were performed in triplicates at least three times.

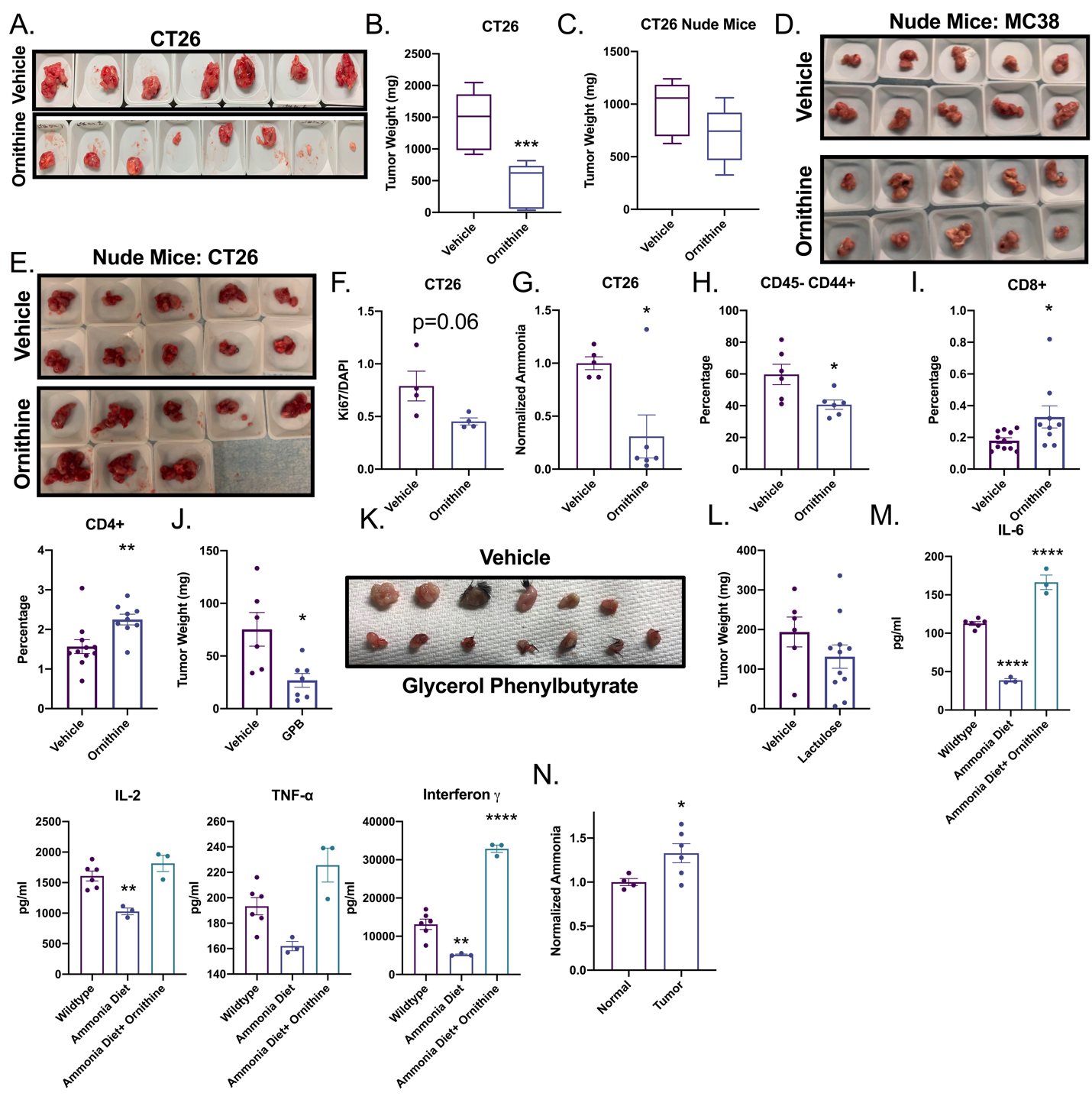

**Figure S6: Ammonia neutralization of colorectal cancer tumor growth is T cell dependent.** A and B) Syngeneic tumor study in BALB/C or C) athymic nude mice with CRC-derived CT26 cells following ornithine treatment (10 mM ornithine intraperitoneally (n=8-10). D and E) Representative images of MC38 and CT26 tumors excised from nude mice. F) Quantification of F) KI67 and G) ammonia in tumors from CT26 (n=3). H and I) T cells numbers in tumors 5 days after ornithine treatment from MC38 implanted syngeneic tumors. (n=4-6). J) Tumor weight and K) images after treatment with glycerol phenylbutyrate daily (n=4). L) Syngeneic tumor study in C57BL/J mice with CRC-derived MC38 cells following lactulose treatment (n=4-5). M) Mice were placed on ammonium acetate diet for 7 days, then CD3 splenic T cells were isolated and plated *ex vivo* for 24 h. The supernatants were collected and an ELISA performed for the indicated cytokine. N) Serum was collected from tumor and normal mice and ammonia was quantified (n=3). *p < 0.05, ^∗∗^ p < 0.01, ^∗∗∗^ p < 0.001, ^∗∗∗∗^ p < 0.0001 unless otherwise indicated. Data is presented as mean +/- the standard error of the mean. All experiments were performed in triplicates at least three times.

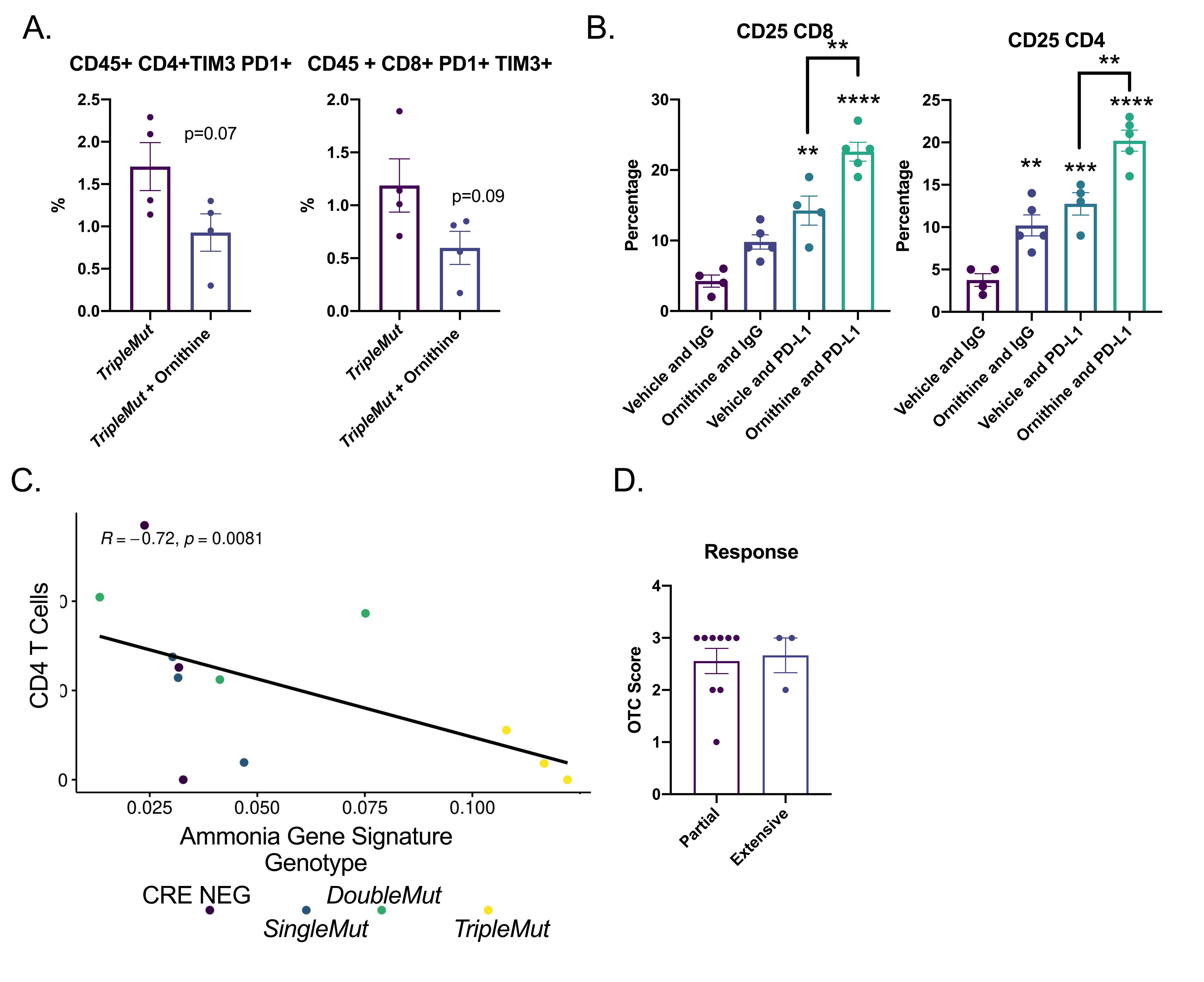

**Figure S7: High ammonia levels are associated with increased T cell exhaustion.** A) Flow populations from colons of induced *TripleMut* mice treated with ornithine and stained for the indicated population (n=3). B) Flow for various cell populations from *TripleMut* mice treated with ornithine, IgG, or anti-PD-L1 therapy for 7 days. (n=3). C) RNA-SEQ samples were evaluated for expression of our ammonia gene signature. Cell populations were deconvoluted using single cell data and compared to the ammonia gene signature (n=3). D) OTC IHC expression grouped into partial or extensive chemotherapy response. Survival significance is calculated using log-rank test. *p < 0.05, ^∗∗^ p < 0.01, ^∗∗∗^ p < 0.001, ^∗∗∗∗^ p < 0.0001 unless otherwise indicated. Data is presented as mean +/- the standard error of the mean. All experiments were performed in triplicates at least three times.

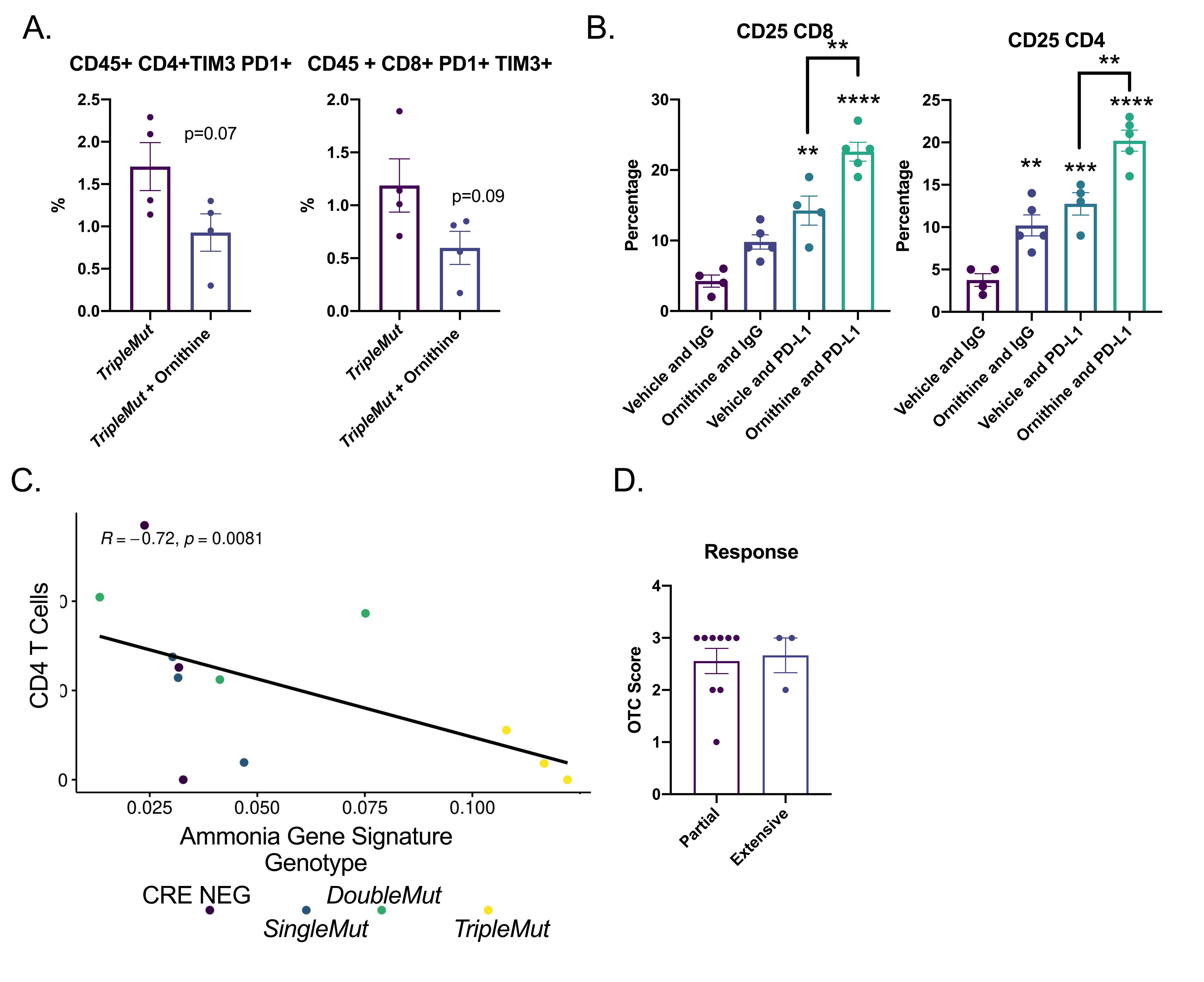

**Figure S7: High ammonia levels are associated with increased T cell exhaustion.** A) Flow populations from colons of induced *TripleMut* mice treated with ornithine and stained for the indicated population (n=3). B) Flow for various cell populations from *TripleMut* mice treated with ornithine, IgG, or anti-PD-L1 therapy for 7 days. (n=3). C) RNA-SEQ samples were evaluated for expression of our ammonia gene signature. Cell populations were deconvoluted using single cell data and compared to the ammonia gene signature (n=3). D) OTC IHC expression grouped into partial or extensive chemotherapy response. Survival significance is calculated using log-rank test. *p < 0.05, ^∗∗^ p < 0.01, ^∗∗∗^ p < 0.001, ^∗∗∗∗^ p < 0.0001 unless otherwise indicated. Data is presented as mean +/- the standard error of the mean. All experiments were performed in triplicates at least three times.
